# North American fireflies host low bacterial diversity

**DOI:** 10.1101/2020.10.06.328070

**Authors:** Emily A. Green, Scott R. Smedley, Jonathan L. Klassen

## Abstract

Although there are numerous studies of firefly mating flashes, lantern bioluminescence, and anti-predation lucibufagin metabolites, almost nothing is known about their microbiome. We therefore used 16S rRNA community amplicon sequencing to characterize the gut and body microbiomes of four North American firefly taxa: *Ellychnia corrusca*, the *Photuris versicolor* species complex, *Pyractomena borealis*, and *Pyropyga decipiens*. These firefly microbiomes all have very low species diversity, often dominated by a single species, and each firefly type has a characteristic microbiome. Although the microbiomes of male and female fireflies did not differ from each other, *Ph. versicolor* gut and body microbiomes did, with their gut microbiomes being enriched in *Pseudomonas* and *Acinetobacter*. *Ellychnia corrusca* egg and adult microbiomes were unique except for a single egg microbiome that shared a community type with *E*. *corrusca* adults, which could suggest microbial transmission from mother to offspring. Mollicutes that had been previously isolated from fireflies were common in our firefly microbiomes. These results set the stage for further research concerning the function and transmission of these bacterial symbionts.

## INTRODUCTION

Beetles (Order: Coleoptera) are one of the most diverse insect groups, containing nearly 400,000 species [1]. The beetles in the family Lampyridae are commonly known by many names: fireflies, lightning bugs, glow worms, lamp-lighters, and night-travelers, all describing the bioluminescent lantern that they use to signal to their mates [2]. Fireflies are found on every continent except Antarctica, living in woods, plains, and marshes. North America is home to over 125 firefly species, including *Ellychnia corrusca*, the *Photuris versicolor* species complex, *Pyractomena borealis* and *Pyropyga decipiens* [2]. These 4 types of fireflies occur sympatrically, ranging from the eastern coast of Canada, south to Florida, and west to the Great Plains [2, 3]. All four of these fireflies are carnivorous or omnivorous as larvae, attacking snails as their prey, and/or eating nectar and plant sap [4]. *Ellychnia corrusca, Pyractomena borealis* and *Pyropyga decipiens* are all closely related to each other, and both *E. corrusca* and *Pyropyga decipiens* are diurnal with no lantern [2]. *Ellychnia corrusca, Pyractomena borealis*, and many other North American fireflies use lucibufagins as a chemical defense. Lucibufagins are cardioactive C-24 steroidal pyrenes and a subclass of the bufadienolides [5–8] that create a disagreeable taste to predators and thereby provide fireflies with protection from bats, birds, and spiders [5]. However, the biosynthetic origin of lucibufagins is not known. At least five genera of fireflies, including the groups in this study, ingest the nectar of the common milkweed plant (*Asclepias syriaca L.*). Although eating is not required for adult firefly survival, cardenolides (a type of toxic steroid) found on milkweed plants are ingested by adult fireflies and may be used to produce lucibufagins [9].

In this study, we focused especially on two fireflies with unique lifestyles: *E. corrusca* and the *Photuris versicolor* species complex. *Ellychnia corrusca* is nicknamed the winter firefly because it has a winter-spring activity cycle, in contrast to the late spring-summer activity cycles of most other fireflies [2, 10]. *Ellychnia corrusca* larvae in New England (North America) extend their larval stage across two years instead of one to ensure that they ingest ample calories, emerging in the late fall of their second year as adults and sexually maturing during the winter to start mating, which lasts until late spring [11]. Normally, adult fireflies do not need to feed because they only survive as adults for 2 to 4 weeks [2, 4]. However, *E. corrusca* firefly adults live for ~9 months, and in Massachusetts USA they ingest interstitial fluid and sap from maple trees to help them survive during the cold winters [11]. The *Photuris versicolor* species complex (hereafter: *Ph. versicolor*), contains > 17 firefly species that are differentiated by their male courtship lantern flashes but that cannot be identified from their morphology with confidence. The *Ph. versicolor* adult life cycle occurs during the summer months (June-August), but unlike other fireflies, *Ph. versicolor* females mimic the mating light display of female *Photinus* fireflies to attract *Photinus* males, earning *Ph. versicolor* the nickname “femme fatale” [12]. Once a male *Photinus* firefly is near, the *Photuris versicolor* female will attack and kill the male, gaining nutrients and lucibufagins that she uses to protect herself, her eggs, and her pupae, which do not themselves produce lucibufagins [5]. Many *E. corrusca* populations complete their life cycles before *Ph. versicolor* becomes seasonally abundant, especially where temperatures are colder. However, *Ph. versicolor* predation of *E. corrusca* can occur, particularly in the Southern USA. *Ph. versicolor* will also attack and eat *E. corrusca* to gain protective lucibufagins in the lab [2, 8], which could give insight into the selective pressures that caused *E. corrusca* to mate in the winter instead of in the summer.

Beetles feed on many different substrates, following herbivorous [13], omnivorous [14], xylophagous [15], detritivorous [16], and carnivorous diets [17]. Beetles that belong to similar taxonomic families but that have different diets have distinct microbial communities [17–19]. Diets lacking in nutrients often cause insects to rely on nutritional symbionts to provide vital nutrients [20], as in dung [21], carabid [14], and carrion beetles [22]. Although there is a vast amount of research studying beetle microbiomes, only a handful of studies consider firefly-associated microbes. In those studies, several Mollicute bacteria were isolated from the fireflies *E. corrusca* and *Ph. versicolor*, but this does not give insight into the composition of their microbiome beyond these strains [23–30]. To fill this gap, we used 16S rRNA community amplicon sequencing to survey the microbiomes of *E. corrusca, Photuris versicolor, Pyractomena borealis*, and *Pyropyga decipiens* fireflies. Our results show that fireflies have simple, species-specific microbiomes, and generate hypotheses about how diet and seasonality may drive firefly microbiome structure and function.

## METHODS

### Sample collection

Live *E. corrusca* and *Photuris versicolor* fireflies were collected by S. Smedley during the winter of 2016-17 and spring/summer of 2017 from residential and camping/forest areas within Vernon, Bolton, and Andover, Connecticut, U.S.A. *Pyractomena borealis* larvae and adults were raised in the laboratory by S. Smedley. No wild *Pyractomena borealis* samples were included in this study. After collection, *E. corrusca*, *Ph. versicolor*, and *Pyractomena borealis* fireflies were stored at −80°C in air-filled vials. *Pyropyga decipiens* fireflies were collected by Lynn Faust in summer 2016 from Ohio and Tennessee and stored in 95% ethanol. Collection dates and sample locations are listed in Supplemental File 1.

### Sample preparation & DNA extraction

All fireflies were surface-sterilized using 3 rounds of a 10 second submersion in 70% ethanol, followed by a 10 second submersion in phosphate-buffered saline (PBS) [31]. Surface-sterilized fireflies were dissected into two separate tissues: a gut sample and the remaining carcass “body” sample. DNA from the tissue dissections were extracted using a bead beating and chloroform-isopropanol protocol as described in [32], except using 0.7 g of 1 mm and 0.3 g of 0.1 mm silica/zirconium beads and 5 cycles of bead-beating & chilling on ice. Negative controls containing only the DNA extraction reagents were processed alongside each batch of tissue samples. The DNA concentration of each extract and negative control was determined using the Qubit dsDNA high-sensitivity assay protocol and a Qubit 3.0 fluorimeter (Invitrogen, Carlsbad California).

### 16S V4 rRNA PCR screen & community amplicon sequencing

DNA samples were PCR amplified using primers 515F and 806R (targeting the bacterial 16S rRNA gene V4 region) to determine the presence of bacterial DNA [33]. Ten nanograms of template DNA was added to 5 μl Green GoTaq Reaction Mix Buffer (Promega, Madison, Wisconsin, USA), 1.25 units of GoTaq DNA Polymerase (Promega, Madison, Wisconsin, USA), 10 μmol of each primer, and 300 ng/μl BSA (New England BioLabs Inc. Ipswitch Massachusetts), to which nuclease free H_2_O was added to a volume of 25 μl. Thermocycling conditions (BioRad, Hercules, California) were: 3 min at 95°C, 30 cycles of 30 sec at 95°C, 30 sec at 50°C, and 60 sec at 72°C, followed by a 5 min cycle at 72°C and then an indefinite hold at 4°C. Gel electrophoresis was used to confirm the expected band size of 300–350 bp.

All samples that had a gel band of the expected size were prepared for community amplicon sequencing of the 16S rRNA gene V4 region using an Illumina MiSeq at the University of Connecticut Microbial Analysis, Resources and Services (MARS) facility. Approximately 30 ng of DNA from each sample was added to a 96-well plate containing 10 μmol each of the forward and reverse Illumina-barcoded versions of primers 515F and 806R, 5 μl AccuPrime buffer (Invitrogen, Carlsbad, California), 50 mM MgSO_4_ (Invitrogen, Carlsbad, California), 300 ng/μl BSA (New England BioLabs Inc. Ipswitch, Massachusetts), a 1μmol spike-in of both non-barcoded primers 515F and 806R, and 1 unit AccuPrime polymerase (Invitrogen, Carlsbad, California), to which nuclease-free H_2_O was added to a volume of 50 μl. Reaction mixes were separated into triplicate reactions (each with a volume of 16.7 μl) in a 384 well plate using an epMotion 5075 liquid handling robot (Eppendorf, Hamburg, Germany). The resulting 384 well plate was then transferred to a thermocycler (Eppendorf, Hamburg, Germany), which used the following conditions: 2 min at 95°C, 30 cycles of 15 sec at 95°C, 60 sec at 55°C, and 60 sec at 68°C, followed by a final extension for 5 min at 68°C and then an indefinite hold at 4°C. After PCR, triplicate reactions were re-pooled using the epMotion and DNA concentrations were quantified using a QIAxcel Advanced capillary electrophoresis system (QIAgen, Hilden, Germany). Samples that had concentrations > 0.5 ng/μl were pooled using equal weights of DNA to create the final sequencing libraries. Libraries were then bead-cleaned using Mag-Bind RXNPure plus beads (OMEGA, Norcross, Georgia) in a 1:0.8 ratio of sequencing library to bead volume. Cleaned library pools were adjusted to a concentration of 1.1 ng/μl ± 0.1 ng/μl, which was confirmed using the Qubit dsDNA high-sensitivity assay on a Qubit 3.0 fluorimeter (Invitrogen, Carlsbad, California). Microbial community sequencing on an Illumina MiSeq (Illumina, San Diego, California) was completed in 2 batches, the first composed of 52 samples and the second composed of 195 samples.

### Post-sequencing & bioinformatic analyses

Our DNA sequencing produced 245 16S rRNA community amplicon sequencing datasets, including 22 negative controls. Reads were analyzed using R v3.5.3 [34] and the dada2 v1.11.1 [35] pipeline for amplicon sequence variants (ASVs) (https://benjjneb.github.io/dada2/tutorial.html, accessed: November 11, 2017). Read counts ranged from 2 to 888,868 (Suppl. File S1). Metadata files for the samples were imported into phyloseq v1.26.1 [36], creating a phyloseq R object that was used for subsequent analyses. Reads that were not classified as belonging to the kingdom Bacteria using the SILVA database v128 were removed [37, 38]. ASVs that matched to mitochondria were then removed separately, because SILVA included them in the kingdom Bacteria. Samples were screened for contamination using the decontam v1.2.1 [39] prevalence protocol with a default threshold value of 0.1. No reads were flagged as contamination, and 17 samples with 0 reads were removed, resulting in 1,675 unique ASVs (Suppl. File S1). Negative control samples were not considered further. All samples in the dataset were then rarefied to 10,000 reads and read counts were converted to relative abundances. The final phyloseq object contained 133 samples from 20 *Photuris versicolor* adults, 4 *Pyropyga decipiens* adults, 7 *Pyractomena borealis* larvae, 1 *Pyractomena borealis* adult, 95 *Ellychnia corrusca* adults, and 6 *E. corrusca* eggs. Despite amplification during the initial PCR screen, of the initial 223 insect samples, 74 samples were removed due to low read counts (53 *E. corrusca* adult samples, 3 *E. corrusca* egg samples, 9 *Photuris versicolor* samples, 2 *Pyropyga decipiens* samples, 4 *Pyractomena borealis* adult samples & 3 *Pyractomena borealis* larvae samples). These low read counts may have been due to: 1) high amounts of host DNA acting as a PCR inhibitor; 2) there being a minimal firefly microbiome, leading to limited template concentrations; or 3) other technical issues such as inefficient PCR amplification using primers that contained the Illumina barcodes compared to our initial PCR screen using non-barcoded primers.

Alpha diversity was measured using the phyloseq ‘plot_richness’ command, and Beta diversity was measured using weighted and unweighted unifrac distance metrics. Weighted and unweighted unifrac distances were calculated, ordinated, and viewed using the ‘distance’, ‘ordinate’, and ‘plot_ordinate’ phyloseq commands, respectively. PERMANOVA statistical tests were calculated using vegan v 2.5-4 [40]. Although weighted unifrac (WUF) and unweighted unifrac (UUF) distances were used for each test, to keep the text concise only one test is listed in the text and the complimentary values are presented in Supplemental Tables S1–S4.

Differences in the relative abundances of taxa in firefly microbiomes were compared using DESeq2 [41], using the Parametric fitType, Wald tests, and an adjusted alpha value of 0.01. A heatmap was constructed for the 26 most abundant genera in the dataset using the gplots v3.03 (https://www.rdocumentation.org/packages/gplots) command ‘heatmap.2’ and using the Euclidean and ward.D distance metrics. To avoid redundancy in the heatmap, ASVs were grouped by genus name using the phyloseq command ‘tax_glom’. Covariance between the top 26 taxa were calculated using SpiecEasi v1.0.7 [42], and the SpiecEasi output file was exported into Cytoscape v3.7.2 [43] using igraph v1.2.4.2 [44] for visualization.

The most abundant Mollicute ASVs from our dataset included four *Mesoplasma* ASVs, two *Spiroplasma* ASVs, and one *Entomoplasma* ASV. A phylogenetic tree was constructed to show the relationships between the 16S rRNA sequences of Mollicutes from this study and those that had been previously isolated from fireflies. Reference Mollicute sequences from taxa belonging to the same genera as our firefly ASVs were selected from the SILVA database, especially those Mollicutes that had been isolated from fireflies and other beetles. Additional Mollicute reference sequences for fireflies that were not represented in the SILVA database were downloaded from NCBI. Sequences were aligned using MUSCLE v3.8.31 [45] and trimmed to the same length. The phylogenetic tree, rooted by the 16S rRNA genes from *Bacillus subtilis* and *Mycoplasma haemominutum*, was calculated using a GTRGAMMAI substitution model and 500 bootstrap replicates in RAxML v8.2.11 [46]. The topology of the tree created using these selected sequences agreed with that constructed using all Mollicute sequences in SILVA, ensuring that taxon selection did not bias our phylogenetic analysis.

The commands used for all analyses is attached as Suppl. File S2. All data are available on NCBI under BioProject PRJNA563849. Raw sequencing reads are deposited in SRA under BioSample numbers SAMN14678004 – SAMN14678257.

## RESULTS

### Firefly microbiomes are typically dominated by single taxa

We characterized the bacterial communities in 133 firefly gut and body dissections (Suppl. File S1) using community amplicon sequencing of the 16S rRNA gene. All firefly microbiomes had low α-diversity (Suppl. Fig. S1). Mean Shannon diversity scores for the firefly microbiomes ranged from 0.70 to 2.52, and alpha diversities of *Photuris versicolor* and *E. corrusca* microbiomes differed from each other, as did those from *Pyropyga decipiens* and *E. corrusca* (Dunn test, p = 0.001 and p = 0.003 respectively). A heatmap of the top 26 bacteria genera found in the microbiomes reflects these low levels of α-diversity, with most firefly microbiomes dominated by a single taxon but with minute amounts of other taxa also present (Fig. 1). Unsupervised clustering of these data grouped *E. corrusca* samples tightly together at the left side of the heat map, while the samples from other species clustered together on the right. Samples from different tissue dissections and sexes did not cluster together for any species of firefly.

**Figure 1).**
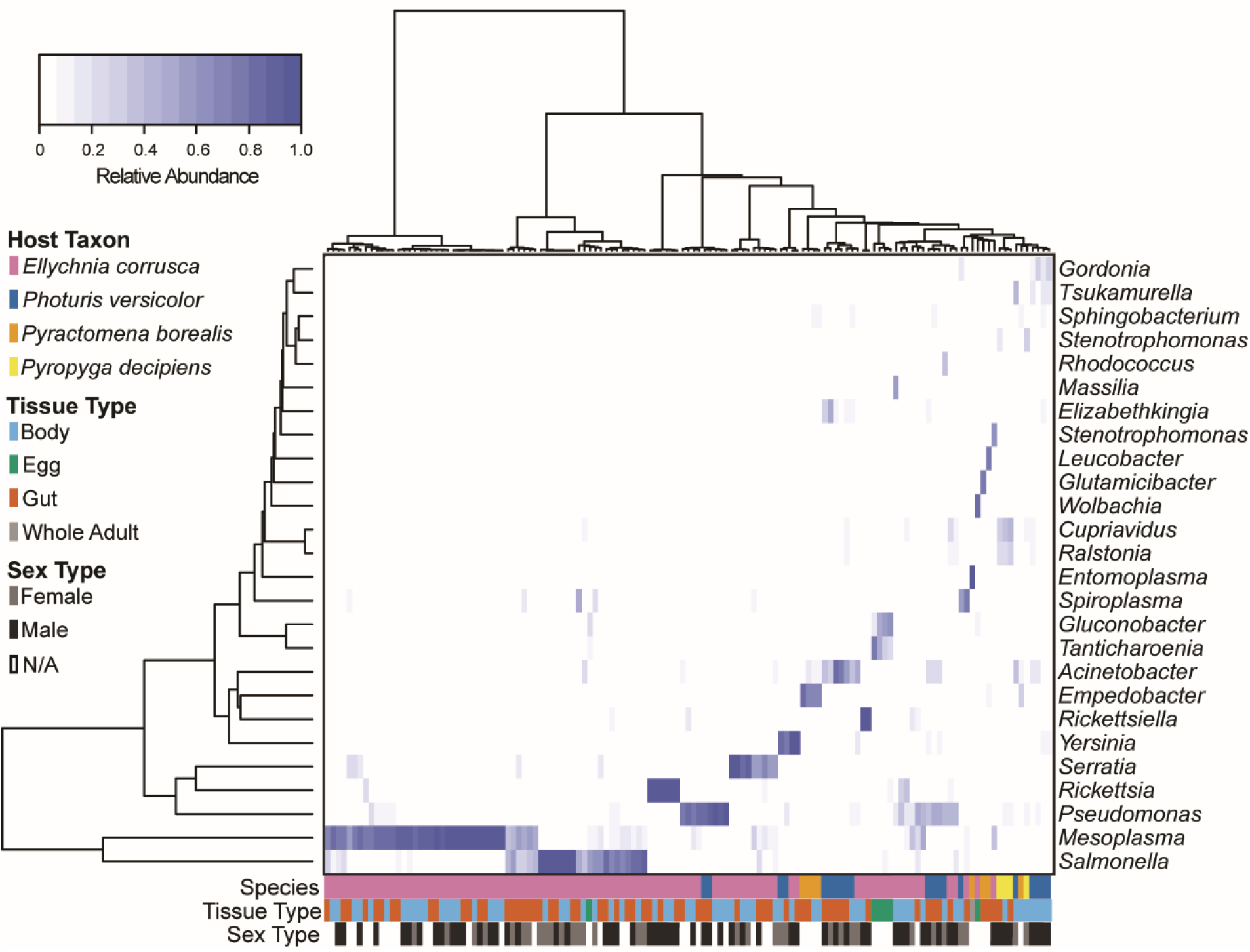
Heat map of the 26 most abundant bacterial genera found in the studied fireflies. To the left, a dendrogram represents the Euclidian distances between the relative abundances of reads assigned to each bacterial genus, which are labeled on the right. The top dendrogram clusters the relative abundances of these genera in each firefly sample using ward.D distances. On the bottom, 3 different rows indicate the host taxon, tissue type, and sex for each sample, indicated by the different colors in the key to the left of the figure. The topmost color key represents the relative abundance of reads in each sample that were assigned to each bacterial genus, with white and dark blue representing 0 and 100% relative abundance, respectively. n = 133

Using these top 26 bacteria genera, we identified taxa whose relative abundances were correlated with each other to infer potential interactions between them (Suppl. Fig. S2). Of the 26 bacteria genera, the relative abundances of 14 genera did not correlate with those of another genus in our dataset, and the relative abundances of 11 genera were positively correlated with those of another genus. The relative abundances of *Ralstonia* and *Cupriavidus*, found in the *Pyropyga decipiens* samples, were strongly and positively correlated with each other, as were the relative abundances of *Tanticharoenia* and *Gluconobacter*, found in *E. corrusca* eggs, and the relative abundances of *Gordonia* and *Tsukamurella*, found in *Photuris versicolor*. Only the relative abundances of *Mesoplasma* and *Acinetobacter* were negatively correlated with each other, and these taxa were not found together in any sample.

In the heatmap, many samples clustered together that were dominated by single taxa. Based on these clusters, we defined community types defined by the genus or genera of bacteria that were present in these samples with relative abundances ≥ 30% (Fig. 2). *Ellychnia corrusca* adults were assigned to 6 distinct community types: T-1 (*Mesoplasma*), T-2 (*Salmonella*), T-3 (*Mesoplasma* & *Salmonella*), T-4 (*Serratia*), T-5 (*Rickettsia*) and T-6 (*Pseudomonas*). *Ellychnia corrusca* egg microbiomes were all assigned to T-13 (*Gluconobacter*), except for a single egg microbiome that belonged to T-2 (*Salmonella*), the same as some *E. corrusca* adult microbiomes. *Photuris versicolor* microbiomes were assigned to 3 community types: T-8 (*Acinetobacter*), T-9 (*Yersinia*) and T-6 (*Pseudomonas*). *Pyractomena borealis* and *Pyropyga decipiens* were assigned to T-10 (*Empedobacter*) and T-12 (*Cupriavidus*), respectively.

**Figure 2).**
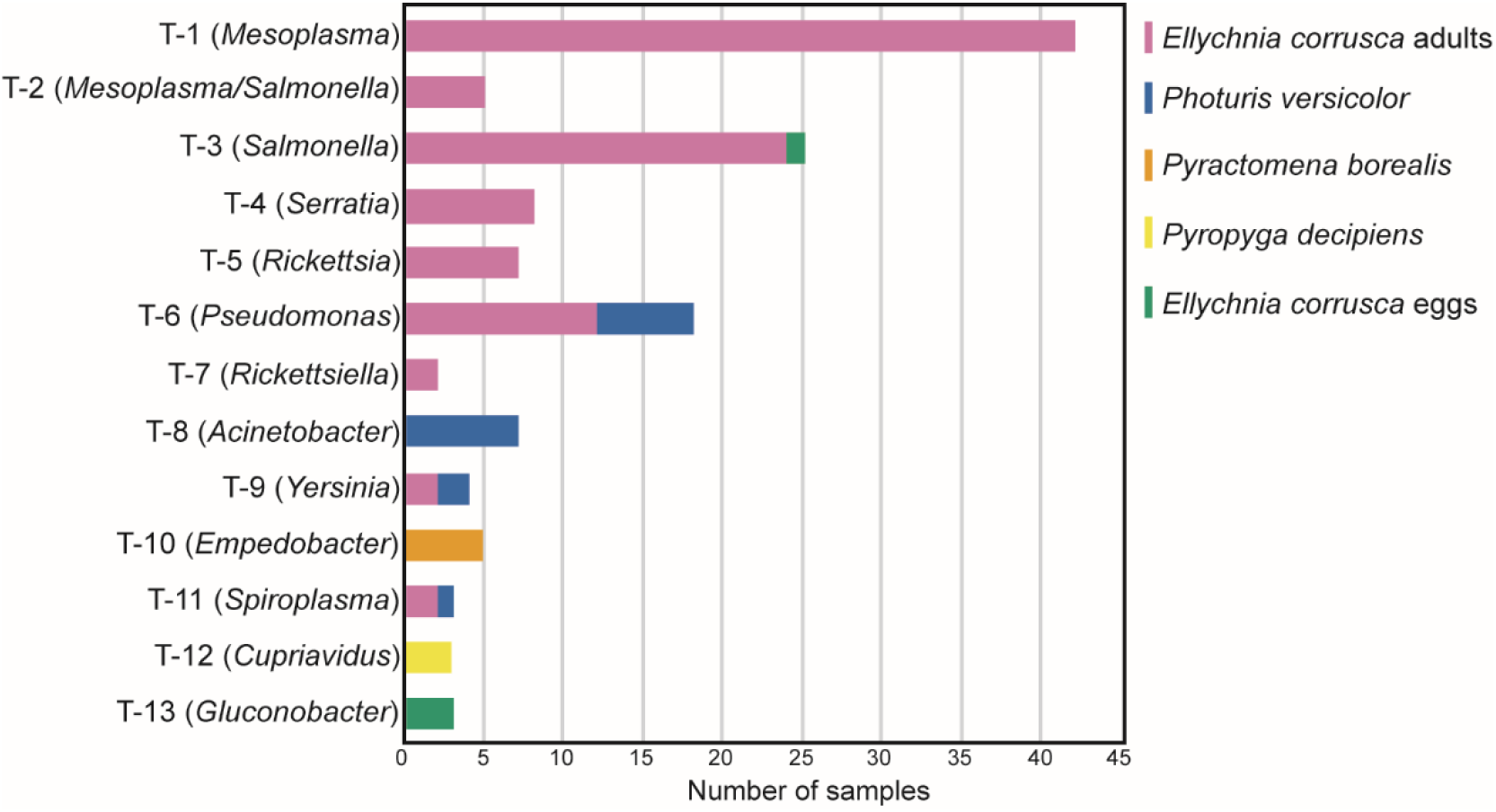
Firefly microbial community types. The X axis indicates the number of samples that were assigned to each community type. Colors indicate firefly taxa, as indicated by the key. Note that some gut and body samples originate from the same individual firefly and so are counted twice.

### Firefly species harbor unique microbial communities

The four sampled firefly taxa had distinct bacterial communities (Weighted Unifrac [WUF] PERMANOVA: R^2^=0.215, p=0.001; Fig. 3A, Suppl. Fig. S3, Suppl. Table S1). In a PCoA ordination of WUF beta-diversity distances, *E*. *corrusca* samples clustered together, spanning from left to right (Fig. 3). *Pyropyga decipiens* samples clustered together with some *E. corrusca* samples and those from *Photuris versicolor* on the left side of Figure 3A, and *Ph versicolor* samples grouped together down the left side and at the bottom, along with the *Pyractomena borealis* samples. Species-specific clustering can be seen in the UUF plot, with *E. corrusca* having the most variance (Suppl. Fig. S3B). The separation between the *E. corrusca* and *Ph versicolor* microbiomes was maintained in a better-balanced comparison when the few *Pyropyga decipiens* and *Pyractomena borealis* samples were excluded (WUF PERMANOVA: R^2^=0.164, p=0.001; Suppl. Fig S4, Suppl. Table S2).

**Figure 3).**
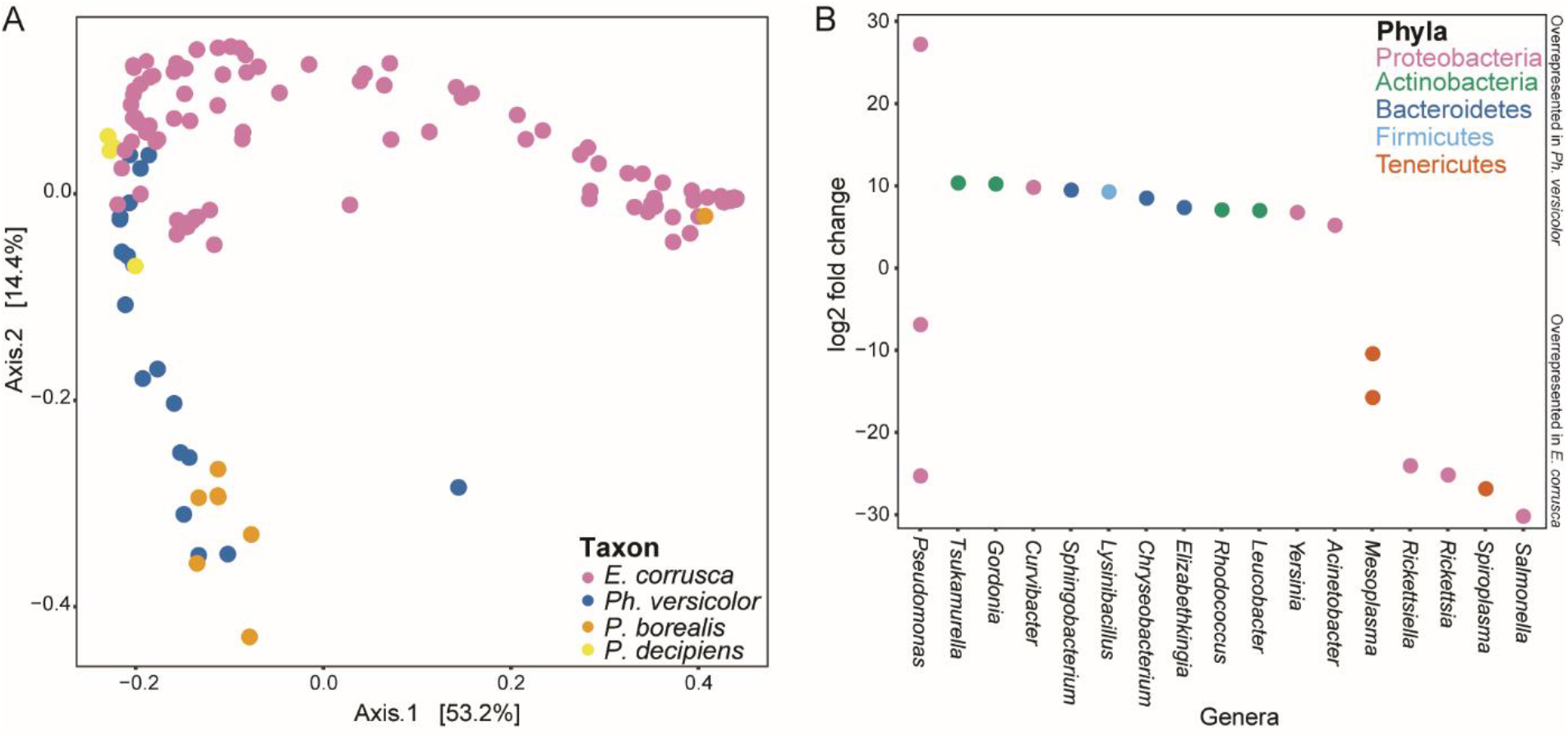
A: PCoA of Weighted Unifrac distances between microbial communities for all four firefly taxa. Species are differentiated using colors. *P. borealis* = *Pyractomena* borealis, *P. decipiens* = *Pyropyga decipiens*. n = 133. B: Over- and underrepresented ASV sequences in *E. corrusca* and *Photuris versicolor*. adults, including both gut and body samples. The X-axis indicates the genera of bacteria whose relative abundances differed between firefly taxa, and the Y-axis indicates the log2 fold change in relative abundance between samples, where the higher numbers indicate overrepresentation in *Ph. versicolor* and negative numbers indicate overrepresentation in *E. corrusca*. Colors indicate phyla. n = 114

We identified the bacterial taxa whose relative abundance differed between *E. corrusca* and *Photuris versicolor* fireflies using DESeq. Although *Pseudomonas* ASVs were found in both fireflies, one *Pseudomonas* ASV was more abundant in *Ph. versicolor* and two other *Pseudomonas* ASVs were more abundant in *E. corrusca* (Fig. 3B). Two *Mesoplasma* ASVs, *Salmonella*, *Serratia*, *Rickettsia* and *Rickettsiella* were all more abundant in *E. corrusca* than in *Ph. versicolor*, and *Tsukamurella*, *Gordonia*, *Curvibacter* and *Sphingobacterium* were all more abundant in *Ph. versicolor* than in *E. corrusca* (Fig. 3B).

### Gut and body microbiomes are distinct, but these differences are firefly-specific

The above analysis indicated that the four sampled firefly groups all hosted distinct microbiomes (Fig. 3). However, further analyses using these same data also indicated that different firefly tissues might also host distinct microbiomes, in a species-specific manner (WUF PERMANOVA for the interactions between species and tissue: R^2^=0.048, p=0.003; Suppl. Fig. S3, Suppl. Table S1). In contrast, a parallel analysis using only samples for which sex was determined indicated that microbiomes did not differ between the sexes, regardless of species or tissue type (all PERMANOVAs testing for differences between sexes: p > 0.05; Suppl. Fig. S5, Suppl. Table S3). In the analysis described above with *Pyractomena borealis* and *Pyropyga decipiens* samples removed to avoid possible artifacts due to unbalanced sample sizes, firefly tissue microbiomes again differed in a species-specific manner (WUF PERMAONVA for interactions between *E. corrusca* and *Photuris versicolor* gut and body microbiomes: R^2^=0.053, p=0.001; Suppl. Fig. S4, Suppl. Table S2).

Because these PERMANOVAs indicated that tissue microbiomes differed in a species-specific manner, we repeated our analysis for each species separately. *Ellychnia corrusca* eggs, gut, and body microbiomes differed from each other (WUF PERMANOVA: R^2^=0.122, p=0.001), with most gut and body samples clustered together in the PCoA and most egg samples clustered separately (Fig. 4A, Suppl. Table S4, Suppl. Fig. S6). When the egg samples were removed from the analysis, *E. corrusca* gut and body microbiomes differed from each other only when using the UUF distance metric (UUF PERMANOVA: R^2^= 0.072, p=0.001; WUF: Suppl. Table S4). *Wolbachia* had low relative abundance in all *E. corrusca* samples, but was the only bacterial genus that was more abundant in *E. corrusca* eggs than in adults. *Mesoplasma*, *Pseudomonas*, *Acinetobacter* and several other genera were all more abundant in *E. corrusca* adults than in eggs (Fig. 4B). *Ellychnia corrusca* eggs and adults also had distinct community types, with *E. corrusca* egg microbiomes mainly assigned to T-13 (*Gluconobacter*), and only a single egg microbiome assigned to T-3 (*Salmonella*) like those of *E. corrusca* adults (Fig. 2). In contrast, *Photuris versicolor* gut and body microbiomes more strongly differed from each other (WUF PERMANOVA: R^2^=0.340, p=0.001; Fig. 4C, Suppl. Fig. S7, Suppl. Table S4), with *Pseudomonas, Acinetobacter, Leucobacter, Elizabethkingia, Curvibacter* and *Achromobacter* all being more abundant in *Ph. versicolor* guts than in bodies (Fig. 4D). *Pyropyga decipiens* gut and body microbiomes did not differ from each other (WUF PERMANOVA: R^2^=0.312, p=0.667; Suppl. Table S4), and *Pyractomena borealis* gut and body microbiomes differed only when using the UUF distance metric (UUF PEMANOVA: R^2^= 0.616, p=0.008; Suppl. Table S4).

**Figure 4).**
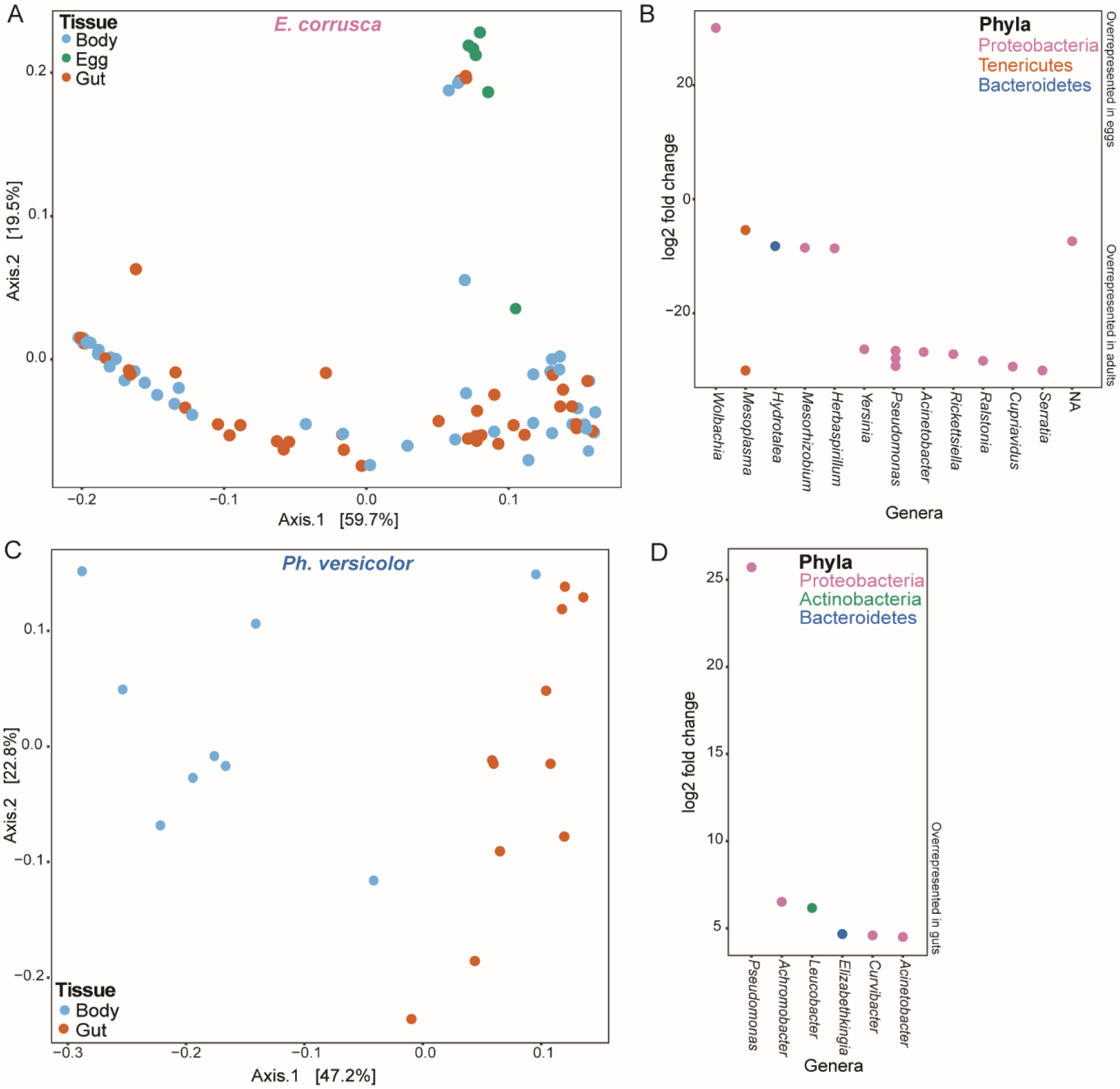
A) PCoA of Weighted Unifrac distances between microbiomes of *E. corrusca* eggs and adults. Egg, gut and body samples are differentiated using colors. n = 100. B) Over- and underrepresented ASV sequences in *E. corrusca* egg and adult samples. The X-axis indicates the genera of bacteria whose relative abundance differed between *E. corrusca* eggs and adults, and the Y-axis indicates the log2 fold change in these relative abundances between tissues, where the higher numbers indicate overrepresentation in eggs and negative numbers indicate overrepresentation in adults. Colors indicate phyla. C) PCoA of Weighted Unifrac distances between microbiomes of *Photuris versicolor* gut and body samples. Gut and body samples are differentiated using colors. D) Over- and underrepresented ASV sequences of *Ph. versicolor* gut and body samples. The X-axis indicates the genera of bacteria whose relative abundance differed between *Ph*. *versicolor* gut and body samples, and the Y-axis indicates the log2 fold change in these relative abundances between tissues, where the higher numbers indicate an overrepresentation in the body samples. Colors indicate phyla. n = 20

These conclusions should be considered preliminary because of the small sample sizes available for both *Pyropaga decipiens* and *Pyractomena borealis*.

### Fireflies host an abundance of potentially symbiotic Mollicutes

Mollicutes were the most abundant bacteria in our 133 firefly samples, consistent with the handful of Mollicutes that had previously been isolated from different firefly species [23–30]. We therefore created a phylogenetic tree to discover how our most abundant Mollicute ASVs were related to the sequences of these known firefly Mollicute isolates (Fig. 5). The most prevalent *Spiroplasma* ASV (ASV Spiroplasma1), found in many of the *E. corrusca* samples, was similar to the 16S rRNA sequence of *S. corruscae*, a species that was first isolated from *E. corrusca* [25]. ASV Spiroplasma2 was found in one *Photuris versicolor* sample, and was similar to the 16S rRNA sequence of *S. ixodetis*, a male-killing agent in butterflies [47]. *S. ixodetis* has not been found in fireflies to date. The four *Mesoplasma* ASVs, mainly found in *E. corrusca*, are all similar to the 16S rRNA sequences for *M. corruscae* and *Entomoplasma ellychniae*, which had both been isolated previously from *E. corrusca* [26, 28, 30]. Although originally separated using cell morphology and culture media requirements, the genera *Mesoplasma* and *Entomoplasma* are paraphyletic [48]. Our phylogenetic tree showing that both *M. corruscae* and *E. ellychniae* cluster tightly together, as seen by Gasparich et al. [48], who suggest that the genus *Mesoplasma* should be changed to *Entomoplasma*. This would mean that *M. corruscae* would be reclassified as a member of the genus *Entomoplasma*, along with all other *Mesoplasma* species. The single *Entomoplasma* ASV, found in *Pyractomena borealis* adults, closely resembled the 16S rRNA sequence for *E. somnilux*, which had been previously isolated from *Pyractomena angulata* [24]. This phylogenetic tree shows that the Mollicutes sequences detected in study are highly similar to strains that had been previously isolated from fireflies, except for that related to *S. ixodetis*.

**Figure 5).**
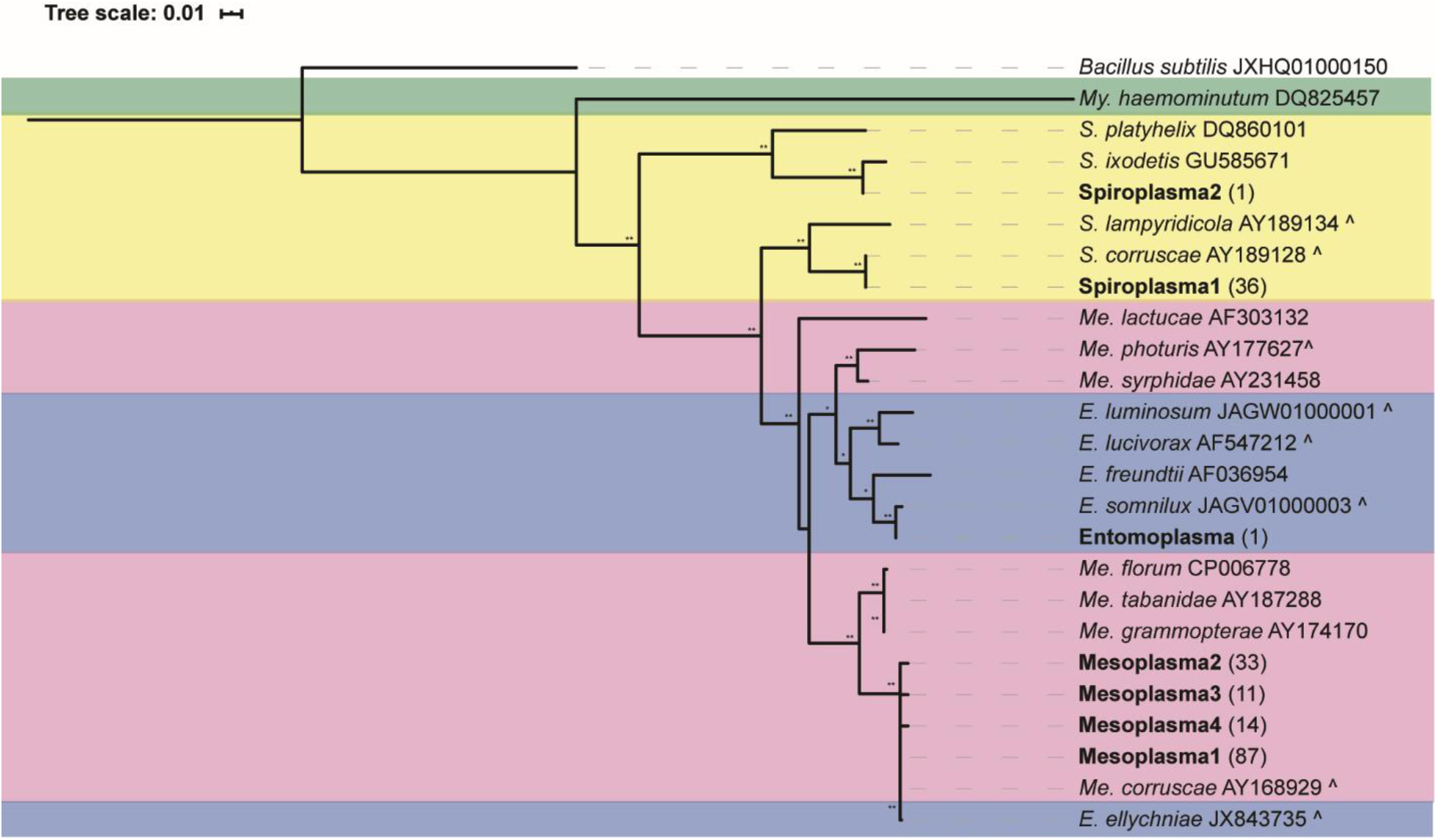
Phylogenetic tree of the Mollicute 16S rRNA gene ASVs in our dataset (in bold) and reference 16S rRNA gene sequences, particularly those isolated from fireflies and beetles (indicated by a ^). The tree was constructed using RAxML with 500 bootstraps and rooted using *Bacillus subtilis* and *Mycoplasma haemominutum*. Bootstrap values of 60–79 and 80–100 are indicated by * and **, respectively. Numbers in parentheses indicate the number of samples in which that ASV was found, and colors group sequences from the same genera. NCBI accession numbers are shown to the right of each reference sequence. *Me* = *Mesoplasma, My* = *Mycoplasma*.

## DISCUSSION

Our results show that all of our sampled fireflies have low-complexity microbiomes. The alpha diversity of our firefly microbiomes was very low, and all samples had only a few taxa present in high abundance (Fig 1). A previous survey also suggested that beetles had low-diversity microbiomes, with carnivorous and herbivorous species hosting the lowest diversity [17]. Our data is consistent with this, as *Photuris versicolor* fireflies are carnivorous and *E. corrusca* ingest plant sap. Kolasa et. al [17]also describe large variation in microbiome composition within a host beetle species, which is consistent with our firefly microbiome data. However, the Shannon diversities measured for our firefly microbiomes (means ranging from 0.7 to 2.52) is lower than those reported in that beetle survey [17]. The low number of correlations between the relative abundances of firefly taxa (Suppl Fig. S2) is consistent with this predominance of simple microbiomes, because (by definition) few interactions can occur when only a single bacterium is abundant in a microbiome, and correlations will inevitably be weak when there is high variability between microbial communities. This variance and low diversity could be explained by microbiomes being acquired via neutral community assembly. Fireflies may be exposed to environmental bacteria by chance, and these bacteria may then multiply and form a stable microbiome. This would explain why there is high variance between the microbiomes from individual insects of the same firefly species.

Samples could be assigned to one of thirteen community types, defined by the taxa in these samples with >30% abundance. Different firefly taxa rarely shared the same community type, and most community types were defined by a single taxon, with only one community type having two taxa that were each >30% abundant (Fig. 2). The presence of multiple community types in each firefly species raises questions about whether the bacteria are either: 1) resident members replicating within the firefly, or 2) transient members that replicate more slowly than the rate of expulsion from the firefly [49]. Although such transient microbiomes are not likely to be conserved between fireflies, it is possible that they do have some effect on their hosts (positive or negative). Community types vary within the same firefly species, suggesting that these microbiomes are transient, but some bacteria were found in extremely high relative abundances (Fig. 1) that could suggest a stable resident microbiome of unknown function. It is possible that each community type provides a different function for the firefly, or that the multiple, highly abundant bacteria in different community types, such as *Mesoplasma*, *Salmonella*, and *Pseudomonas*, could provide similar functions for their hosts. However, this may be unlikely for Mollicute bacteria due to their reduced genomes.

Fireflies from different taxa also had distinct microbiomes (Fig. 3A). *E. corrusca* and *Photuris versicolor* had distinct bacterial communities, and samples from the same species clustered together in PCoA plots of beta-diversity (Suppl. Fig. 4). Differences between the microbiomes of these two groups might be due to their different lifestyles. North American *E. corrusca* are active in winter and may feed on tree sap [10, 11], and so *E. corrusca* microbial communities may therefore be acquired via the ingestion of these fluids. *Ellychnia corrusca* also live for ~9 months. This much longer life span compared to other the firefly taxa in this study could explain the greater diversity of community types in *E. corrusca*. Unlike *E. corrusca, Ph. versicolor* females are predatory and active in the summer. Insect gut microbiomes often vary based on differences in their host’s habitat and diet (e.g., omnivory vs. carnivory) [18], and such differences might underlie the differences that we observed between *E. corrusca* and *Ph. versicolor* microbiomes. Both *E. corrusca* and *Ph. versicolor* hosted *Pseudomonas* ASVs that were similar to those previously found on trees and in soils, which could be explained by *E. corrusca* and *Ph. versicolor* living in soil, on tree bark, and on plant leaves [2], where *Pseudomonas* is common. *Pseudomonas* is also commonly found in beetles from the family Staphylinidae, which are primarily predatory, but not in other non-predatory beetles [17]. This could suggest a relationship between diet and microbiome composition.

All of the *E. corrusca* and *Ph. versicolor* samples used in this study were collected in Connecticut, USA, and all *Pyractomena borealis* samples were-lab raised. *Pyropyga decipiens* samples were collected ~184 miles apart, but had similar microbiome diversities compared to fireflies collected in Connecticut, which suggests that *Pyropyga decipiens* microbiomes may not differ between geographic locations. Whether the microbiomes of *E. corrusca, Ph. versicolor* and *Pyractomena borealis* vary in other geographic locations needs future research.

For some firefly taxa, eggs, guts, and bodies had distinct microbiomes. *Ellychnia corrusca* adult and egg microbiomes differed from each other, with samples clustering separately in PCoAs (Fig. 4A, Suppl. Fig. S6), but an *E. corrusca* egg also had the same community type (T-2; *Salmonella*) as some adults. Although it is unknown which firefly laid this egg, these results suggest that it might be possible for *Salmonella* to be transmitted vertically between *E. corrusca* adults and eggs. The mechanisms that *E. corrusca* may use to acquire these microbes from the environment are unknown, but there are other insects that rely on horizontal transmission for their symbionts, such as the bean bug *Riptortus pedestris*, which acquires its *Burkholderia* symbiont from soil each generation using a mucus-filled organ that the *Burkholderia* specifically penetrates and colonizes [50]. Although *E. corrusca* is not known to have an organ that selects for a specific bacterial symbiont, *Mesoplasma* was found in many samples of *E. corrusca*. It is possible that *E. corrusca* may select this bacterium from the environment, or that *Mesoplasma* selects *E. corrusca* as a preferred host. *E. corrusca* is the most sampled firefly species in our study, and hosts microbiomes belonging to the most community types. This increased sampling may therefore have increased the number of community types detected in this organism relative to the other species in our study.

The microbiomes of the male and female *Photuris versicolor* did not differ from each other, suggesting that the microbiome is not acquired from the firefly prey because only female *Ph. versicolor* are carnivorous, although adult *Ph. versicolor* males do eat milkweed nectar. *Photuris versicolor* gut and body microbiomes also differed from each other, with gut microbiomes having higher amounts of *Pseudomonas* and *Acinetobacter* than body microbiomes (Figs. 4C and 4D). These ASVs may be acquired from the firefly’s environment, because both ASVs were similar to other strains of bacteria that are common in soil and on plants. The bacteria present in the *Ph. versicolor* gut may be derived from eating other fireflies or nectar, and thus the *Pseudomonas* ASVs that are common in *Ph. versicolor* guts may originate from their last meal. The *Ph. versicolor* microbiome may therefore be transient and assembled from the diet microbiome. Alternatively, it is possible that the *Ph. versicolor* gut microbiome is not transient, and the gut provides a niche that is favorable for the growth of *Pseudomonas* and *Acinetobacter* microbes that might originate from soil, plants, or the diet microbiome and subsequently colonize the *Ph. versicolor* gut.

The origin of the body microbiome remains unclear. Body dissections include all other parts of the firefly except for the gut, meaning that other non-gut tissues and hemolymph from inside of the firefly could have been colonized with bacteria. Alternatively, adult firefly abdomens are surrounded with sclerotized segments called ventrites and tergites on their ventral and dorsal sides, respectively [2]. These ventrites and tergites have small gaps between them that allow for movement. Although our firefly samples were ethanol-washed before dissection, this sterilization may not be absolutely perfect, and the body microbiome may therefore include bacteria that are trapped within these segments [51] that were acquired during movement on tree bark or in the leaf litter, where many bacteria are found.

Mollicutes have been isolated from several different firefly species, but their prevalence and function is unclear [23–30]. We detected 4 *Mesoplasma* ASVs in North American fireflies, mainly in *E. corrusca*, that all closely resembled *M. corruscae* and *Entomoplasma ellychniae*, which were both isolated previously from *E. corrusca*. As mentioned in the results, the tight clustering of *M. corruscae* and *Entomoplasma ellychniae* in Figure 5 implies that both belong to the same genus. Gasparich et al. [48] proposed reassigning all *Mesoplasma* species to the genus *Entomoplasma*, including changing *Mesoplasma corruscae* to *Entomoplasma corruscae*. Members of the currently defined genus *Mesoplasma* frequently occur on plants without causing disease, and therefore it could be horizontally acquired by *E. corrusca* when they feed on milkweed nectar or nutrient-poor tree sap to aid in survival during cold winter months [10, 11]. All the fireflies in this study are omnivorous or carnivorous predators as larvae. Other than adult female *Ph. versicolor* species, adult fireflies do not need to eat to survive, although some fireflies might acquire bacteria while feeding as adults. *Pyractomena borealis* larvae are known to ingest tree sap, just as *E. corrusca* larvae do [4], and were colonized by *Entomoplasma somnilux*. Although *Entomoplasma somnilux* has only been found in fireflies, it is possible that it was acquired via the *Pyractomena borealis* diet. Mollicutes and other bacteria may be acquired by a larval firefly’s diet of tree sap and remain colonized through the adult stage. However, it is unknown whether bacteria survive firefly metamorphosis from larvae to adults. If not, the presence or absence of Mollicute in these adult fireflies might be explained by the presence of their diet. Future work will be required to determine the function, if any, of these Mollicutes in their firefly hosts.

Our results provide the first description of the microbiomes found in *E. corrusca*, *Photuris versicolor*, *Pyropyga decipiens* and *Pyractomena borealis*. These microbiomes have low alpha diversities and are species-specific. Some firefly species also have distinct egg, gut, and body microbiomes. Future research will determine the function of these microbiomes and how they are acquired from their environment or transmitted between hosts.

## Supporting information

Supplemental File1: Metadata

Supplemental File2: Bioinformatic Code

## ETHICS APPROVAL

Not applicable

## CONSENT TO PARTICIPATE

Not applicable

## CONSENT FOR PUBLICATION

Emily A. Green and Jonathan L. Klassen approve this for publication. Scott R. Smedley is deceased, and therefore was unable to provide explicit consent.

## AVAILABILITY OF DATA AND MATERIAL

All data are available on NCBI under BioProject PRJNA563849. Raw sequencing reads are deposited in SRA under BioSample numbers SAMN14678004 – SAMN14678257.

## CODE AVAILABILITY

The commands used for all analyses is attached as Suppl. File S2.

## AUTHORS CONTRIBUTIONS

Sample collection: SRS; experiment design: EAG, SRS, and JLK; data analysis: EAG; figure and table creation: EAG; writing (draft preparation and editing): EAG and JLK.

## ACKNOWLEDGEMENTS

We would like to thank Erin L. Mostoller and Dr. Craig W Schneider both from Trinity College, Hartford, CT, and Dr. Steven Deyrup from Siena College, Loudonville, NY for their assistance with obtaining the firefly samples. We would also like to thank the members of the Klassen Lab for their thoughtful feedback on this manuscript before submission, and the UConn Microbial Analysis, Resources, and Services facility for microbiome sequencing.

## WORK CITED

1. Slipinski SA, Leschen RAB, Lawrence JF (2011) Order Coleoptera Linnaeus, 1758. In: Zhang, Z.-Q. (Ed.) Animal biodiversity: An outline of higher-level classification and survey of taxonomic richnes. Zootaxa 3148:203–208. https://doi.org/10.11646/zootaxa.3148.1.26

2. Faust LF (2017) Fireflies, glow-worms, and lightning bugs!: Identification and natural history of the fireflies of the Eastern and Central United States and Canada. University of Georgia Press, Athens, Georgia

3. Barber HS (1951) Fireflies of the genus *Photuris*. Smithson Misc Collect 117:1–66

4. Faust LF, Hughes LS, Zloba MH, Farrington HL (2019) Life history and updated range extension of *Photinus scintillans* (Coleoptera: Lampyridae) with new Ohio records and regional observations for several firefly species. Ohio Biol Surv Notes 9:16–34

5. Eisner T, Goetz MA, Hill DE, et al (1997) Firefly “femmes fatales” acquire defensive steroids (lucibufagins) from their firefly prey. Proc Natl Acad Sci USA 94:9723–9728. https://doi.org/10.1073/pnas.94.18.9723

6. Goetz MA, Meinwald J, Eisner T (1981) Lucibufagins, IV. New defensive steroids and a pterin from the firefly, *Photinus pyralis* (Coleoptera: Lampyridae). Experientia 37:679–680. https://doi.org/10.1007/BF01967916

7. Smedley SR, Risteen RG, Tonyai KK, et al (2017) Bufadienolides (lucibufagins) from an ecologically aberrant firefly (*Ellychnia corrusca*). Chemoecology 27:141–153. https://doi.org/10.1007/s00049-017-0240-6

8. Deyrup ST, Risteen RG, Tonyai KK, et al (2017) Escape into winter: Does a phenological Shift by *Ellychnia corrusca* (winter firefly) shield it from a specialist predator (*Photuris*)? Northeast Nat 24:147–166. https://doi.org/10.1656/045.024.s717

9. Faust L, Faust H (2014) The occurrence and behaviors of North American fireflies (Coleoptera: Lampyridae) on milkweed, *Asclepias syriaca* L. Coleopt Bull 68:283–291. https://doi.org/10.1649/0010-065x-68.2.283

10. Rooney JA, Lewis SM (2000) Notes on the life history and mating behavior of *Ellychnia corrusca* (Coloeptera: Lampyridae). Florida Entomol 83:324–334. https://doi.org/10.2307/3496351

11. Faust L (2012) Fireflies in the snow: Observations on two early-season arboreal fireflies *Ellychnia corrusca* and *Pyractomena borealis*. Lampyrid 2:48–71

12. Lloyd JE (1965) Aggressive Mimicry in *Photuris*: Signal repertoires by femmes fatales. Am Assoc Adv Sci 149:653–654

13. Arias-Cordero E, Ping L, Reichwald K, et al (2012) Comparative evaluation of the gut microbiota associated with the below- and above-ground life stages (larvae and beetles) of the forest cockchafer, *Melolontha hippocastani*. PLoS One 7:e51557. https://doi.org/10.1371/journal.pone.0051557

14. Lundgren JG, Lehman RM (2010) Bacterial gut symbionts contribute to seed digestion in an omnivorous beetle. PLoS One 5:e10831. https://doi.org/10.1371/journal.pone.0010831

15. Grünwald S, Pilhofer M, Höll W (2010) Microbial associations in gut systems of wood- and bark-inhabiting longhorned beetles [Coleoptera: Cerambycidae]. Syst Appl Microbiol 33:25–34. https://doi.org/10.1016/j.syapm.2009.10.002

16. Kaltenpoth M, Steiger S (2014) Unearthing carrion beetles’ microbiome: characterization of bacterial and fungal hindgut communities across the Silphidae. Mol Ecol 23:1251–1267. https://doi.org/10.1111/mec.12469

17. Kolasa M, Ścibior R, Mazur MA, et al (2019) How hosts taxonomy, trophy, and endosymbionts shape microbiome diversity in beetles. Microb Ecol 78:995–1013. https://doi.org/10.1007/s00248-019-01358-y

18. Yun JH, Roh SW, Whon TW, et al (2014) Insect gut bacterial diversity determined by environmental habitat, diet, developmental dtage, and phylogeny of host. Appl Environ Microbiol 80:5254–5264. https://doi.org/10.1128/AEM.01226-14

19. Colman DR, Toolson EC, Takacs-Vesbach CD (2012) Do diet and taxonomy influence insect gut bacterial communities? Mol Ecol 21:5124–5137. https://doi.org/10.1111/j.1365-294X.2012.05752.x

20. Engel P, Moran NA (2013) The gut microbiota of insects - diversity in structure and function. FEMS Microbiol Rev 37:699–735. https://doi.org/10.1111/1574-6976.12025

21. Estes AM, Hearn DJ, Snell-Rood EC, et al (2013) Brood ball-mediated transmission of microbiome members in the dung beetle, *Onthophagus taurus* (Coleoptera: Scarabaeidae). PLoS One 8:e79061. https://doi.org/10.1371/journal.pone.0079061

22. Shukla SP, Plata C, Reichelt M, et al (2018) Microbiome-assisted carrion preservation aids larval development in a burying beetle. Proc Natl Acad Sci U S A 115:11274–11279. https://doi.org/10.1073/pnas.1812808115

23. Hackett KJ, Whitcomb RF, Tully JG, et al (1992) Lampyridae (Coleoptera): A plethora of Mollicute associations. Microb Ecol 23:181–193

24. Williamson DL, Tully JG, Rose DL, et al (1990) *Mycoplasma somnilux* sp. nov., *Mycoplasma luminosum* sp. nov., and *Mycoplasma lucivorax* sp. nov., new sterol-requiring mollicutes from firefly beetles (Coleoptera: Lampyridae). Int J Syst Bacteriol 40:160–164. https://doi.org/10.1099/00207713-40-2-160

25. Hackett KJ, Whitcomb RF, French FE, et al (1996) *Spiroplasma corruscae* sp. nov., from a firefly beetle (Coleoptera: Lampyridae) and tabanid flies (Diptera: Tabanidae). Int J Syst Bacteriol 46:947–950. https://doi.org/10.1099/00207713-46-4-947

26. Tully JG, Rose DL, Hackett KJ, et al (1989) *Mycoplasma ellychniae* sp. nov., a sterol-requiring Mollicute from the firefly beetle *Ellychnia corrusca*. Int J Syst Bacteriol 39:284–289. https://doi.org/10.1099/00207713-39-3-284

27. Stevens C, Tang AY, Jenkins E, et al (1997) *Spiroplasma lampyridicola* sp. nov., from the firefly beetle *Photuris pennsylvanicus*. Int J Syst Bacteriol 47:709–712. https://doi.org/10.1099/00207713-47-3-709

28. Tully JG, Whitcomb RF, Hackett KJ, et al (1994) Taxonomic descriptions of eight new non-sterol-requiring mollicutes assigned to the genus *Mesoplasma*. Int J Syst Bacteriol 44:685–693. https://doi.org/10.1099/00207713-44-4-685

29. Wedincamp J, French FE, Whitcomb RF, Henegar RB (1996) Spiroplasmas and Entomoplasmas (Procaryotae: Mollicutes) associated with tabanids (Diptera: Tabanidae) and fireflies (Coleoptera: Lampyridae). J Invertebr Pathol 68:183–186

30. Tully JG, Bove JM, Laigret F, Whitcomb RF (1993) Revised taxonomy of the class Mollicutes: Proposed elevation of a monophyletic cluster of arthropod-associated Mollicutes to ordinal rank (Entomoplasmatales ord. nov.), with provision for familial rank to separate species with nonhelical morphology. Int J Syst Bacteriol 43:630–630. https://doi.org/10.1099/00207713-43-3-630a

31. Sanders JG, Powell S, Kronauer DJC, et al (2014) Stability and phylogenetic correlation in gut microbiota: Lessons from ants and apes. Mol Ecol 23:1268–1283. https://doi.org/10.1111/mec.12611

32. Lee KM, Adams M, Klassen JL (2019) Evaluation of DESS as a storage medium for microbial community analysis. PeerJ 7:e6414. https://doi.org/10.7717/peerj.6414

33. Caporaso JG, Lauber CL, Walters WA, et al (2011) Global patterns of 16S rRNA diversity at a depth of millions of sequences per sample. Proc Natl Acad Sci U S A 108:4516–4522. https://doi.org/10.1073/pnas.1000080107

34. R Core Team (2018) R: A Language and Environment for Statistical Computing

35. Callahan BJ, McMurdie PJ, Rosen MJ, et al (2016) DADA2: High-resolution sample inference from Illumina amplicon data. Nat Methods 13:581–583. https://doi.org/10.1038/nmeth.3869

36. McMurdie PJ, Holmes S (2013) Phyloseq: An R package for reproducible interactive analysis and graphics of microbiome census data. PLoS One 8:e61217. https://doi.org/10.1371/journal.pone.0061217

37. Quast C, Pruesse E, Yilmaz P, et al (2013) The SILVA ribosomal RNA gene database project: Improved data processing and web-based tools. Nucleic Acids Res 41:590–596. https://doi.org/10.1093/nar/gks1219

38. Yilmaz P, Parfrey LW, Yarza P, et al (2014) The SILVA and “all-species Living Tree Project (LTP)” taxonomic frameworks. Nucleic Acids Res 42:643–648. https://doi.org/10.1093/nar/gkt1209

39. Davis NM, Proctor DiM, Holmes SP, et al (2018) Simple statistical identification and removal of contaminant sequences in marker-gene and metagenomics data. Microbiome 6:226. https://doi.org/10.1186/s40168-018-0605-2

40. Oksanen J, Blanchet FG, Friendly M, et al (2019) vegan: Community Ecology Package. R package version 2.5-4

41. Love MI, Huber W, Anders S (2014) Moderated estimation of fold change and dispersion for RNA-seq data with DESeq2. Genome Biol 15:e550. https://doi.org/10.1186/s13059-014-0550-8

42. Kurtz ZD, Müller CL, Miraldi ER, et al (2015) Sparse and compositionally robust inference of microbial ecological networks. PLOS Comput Biol 11:e1004226. https://doi.org/10.1371/journal.pcbi.1004226

43. Shannon P, Markiel A, Ozier O, et al (2003) Cytoscape: a software environment for integrated models of biomolecular interaction networks. Genome Res 13:2498–2504. https://doi.org/10.1101/gr.1239303

44. Csardi G, Nepusz T (2005) The igraph software package for complex network research. InterJournal Complex Syst 1695

45. Edgar RC (2004) MUSCLE: multiple sequence alignment with high accuracy and high throughput. Nucleic Acids Res 32:1792–1797. https://doi.org/10.1093/nar/gkh340

46. Stamatakis A (2014) RAxML version 8: a tool for phylogenetic analysis and post-analysis of large phylogenies. Bioinformatics 30:1312–1313. https://doi.org/10.1093/bioinformatics/btu033

47. Binetruy F, Bailly X, Chevillon C, et al (2019) Phylogenetics of the *Spiroplasma ixodetis* endosymbiont reveals past transfers between ticks and other arthropods. Ticks Tick Borne Dis 10:575–584. https://doi.org/10.1016/j.ttbdis.2019.02.001

48. Gasparich GE, Kuo C (2019) Genome analysis-based union of the genus *Mesoplasma* with the genus *Entomoplasma*. Int J Syst Evol Microbiol 69:2735–2738. https://doi.org/10.1099/ijsem.0.003548

49. Hammer TJ, Sanders JG, Fierer N (2019) Not all animals need a microbiome. FEMS Microbiol Lett 366:fnz117. https://doi.org/10.1093/femsle/fnz117

50. Ohbayashi T, Takeshita K, Kitagawa W, et al (2015) Insect’s intestinal organ for symbiont sorting. Proc Natl Acad Sci U S A 112:5179–5188. https://doi.org/10.1073/pnas.1511454112

51. Hammer TJ, Dickerson JC, Fierer N (2015) Evidence-based recommendations on storing and handling specimens for analyses of insect microbiota. PeerJ 8:e1190. https://doi.org/10.7717/peerj.1190

