## Supplemental File3: Supplemental Figures and tables for "North American fireflies host low bacterial diversity"

^c^ Deceased

Emily A Green: https://orcid.org/0000-0002-0824-6369

Jonathan Klassen PhD: https://orcid.org/0000-0003-1745-8838

Supplementary Figures and Tables

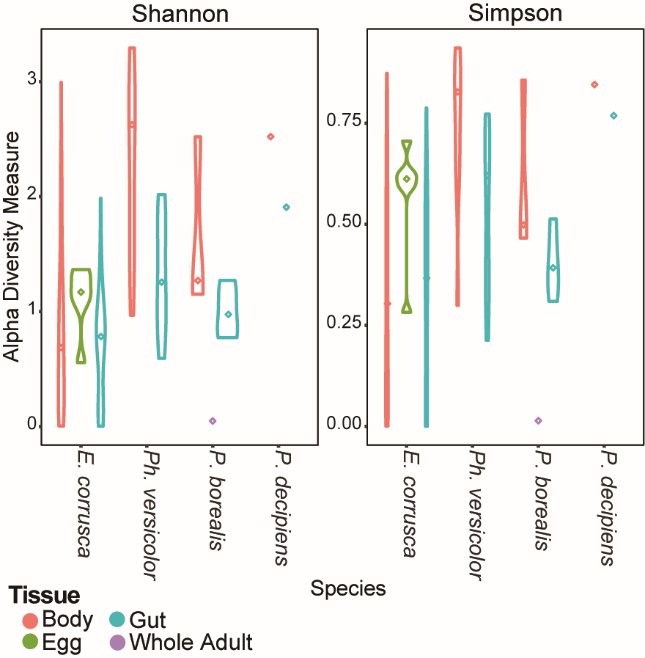

Supplemental Figure S1) Shannon and Simpson alpha-diversities for all four sampled firefly taxa, grouped by body, egg, and gut samples. The X-axes list each firefly taxon, and the median α-diversity is marked with a diamond. *P. borealis* = *Pyractomena borealis*, *P. decipiens* = *Pyropyga decipiens*. Colors indicate tissue type. n = 133

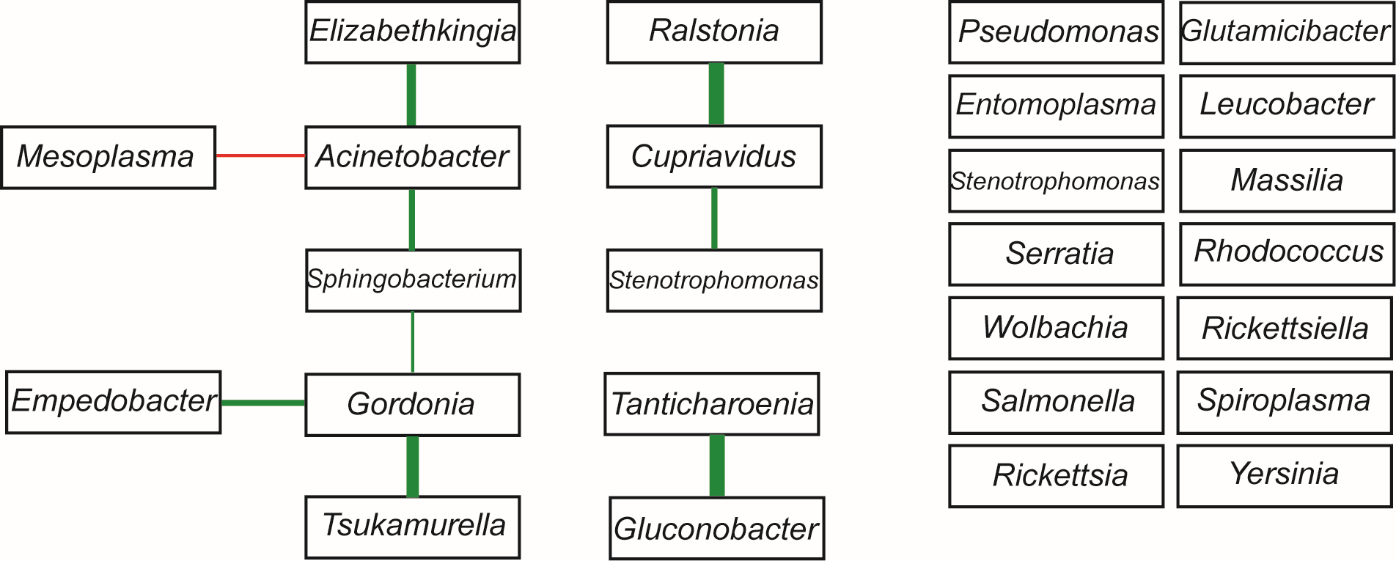

Supplemental Figure S2) Spiec-Easi correlation matrix for the top 26 genera found in *E. corrusca*, *Ph. versicolor*, *Pyropyga decipiens*, and *Pyractomena borealis* microbiomes. Positive correlations between genera are indicated using green lines, and negative correlations by red lines. Genera whose relative abundances did not correlate to each other lack connecting lines. The line thickness is proportional to correlation strength.

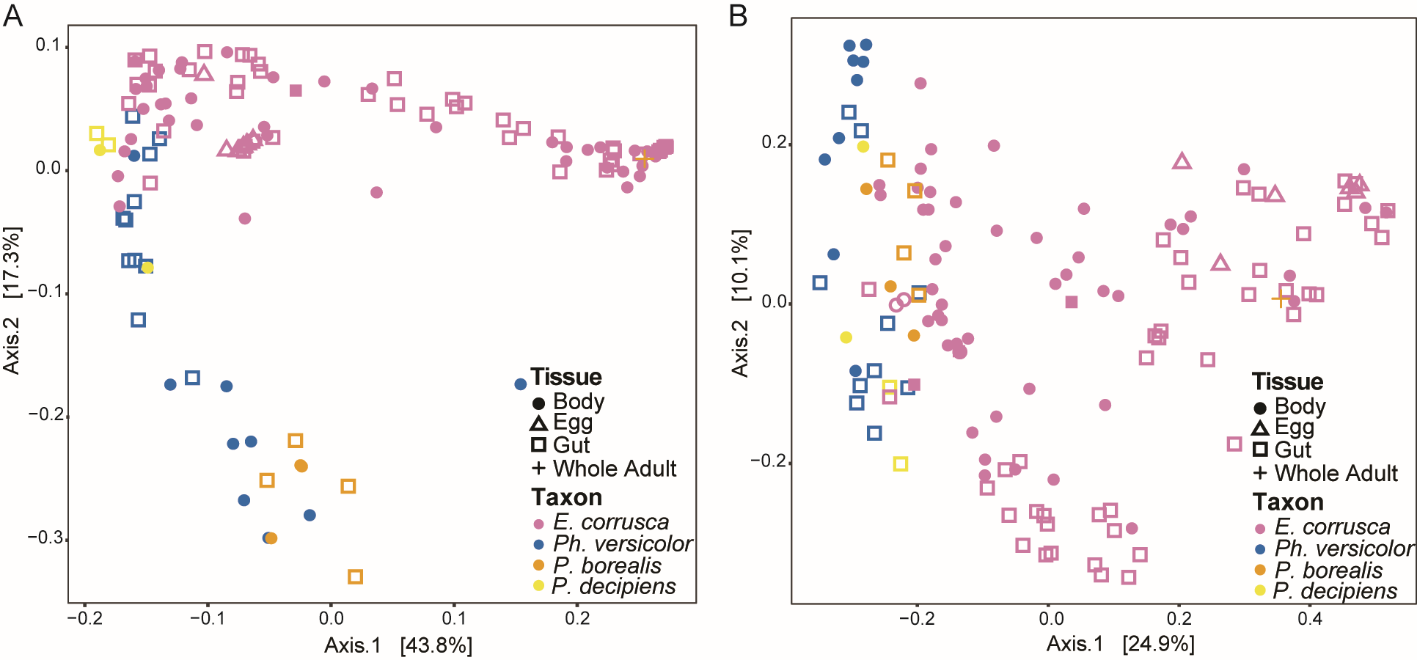

Supplemental Figure S3) PCoA plot of weighted (A) and unweighted (B) unifrac distances between microbiomes for all four sampled firefly taxa and tissue types. Gut, body, egg and whole adult samples are indicated by shape, and taxa are indicated by color. *P. borealis* = *Pyractomena* borealis, *P. decipiens* = *Pyropyga decipiens*. n = 133

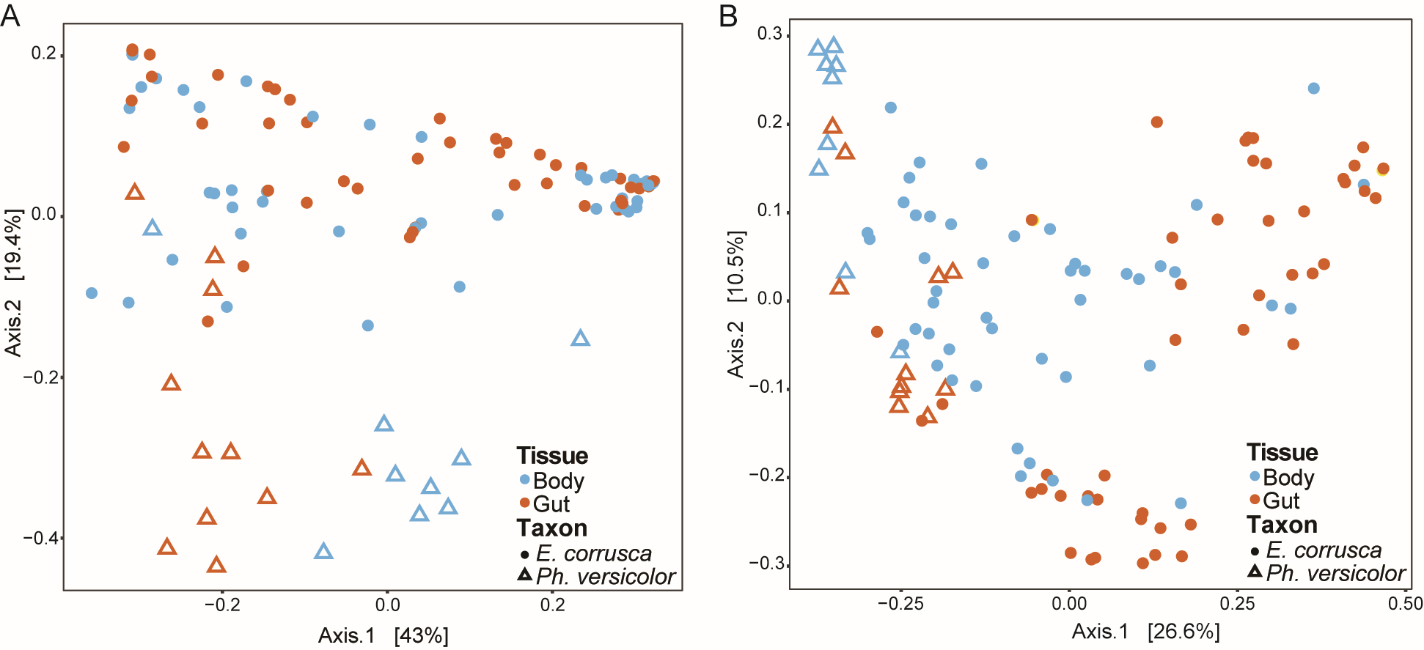

Supplemental Figure S4) PCoA plots of weighted (A) and unweighted (B) unifrac distances between microbiomes from *E. corrusca* and *Ph. versicolor* adults. Gut and body samples are indicated by shape, and taxa are indicated by color. n = 114

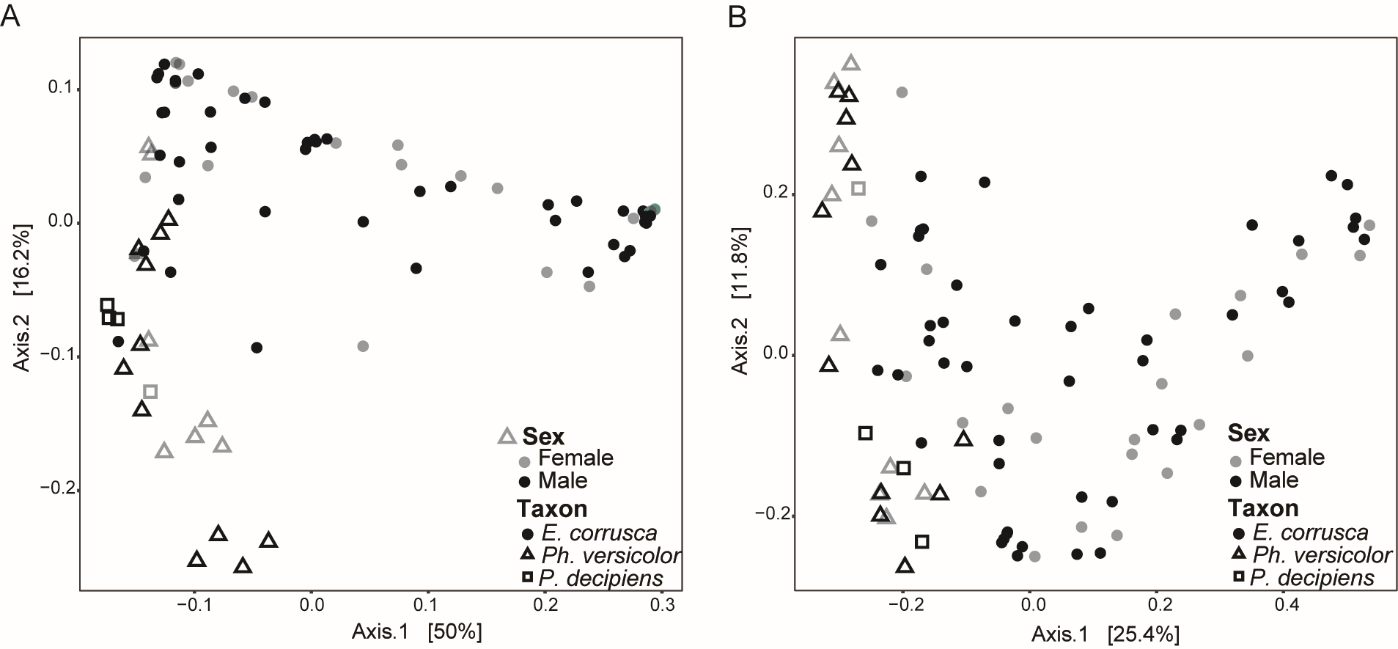

Supplemental Figure S5) PCoA plots of weighted (A) and unweighted (B) unifrac distances between microbiomes from *E. corrusca*, *Ph. versicolor* and *Pyropyga decipiens* adults that had an identified sex. Sexes are indicated by shade, and taxa are indicated by shape. n = 92

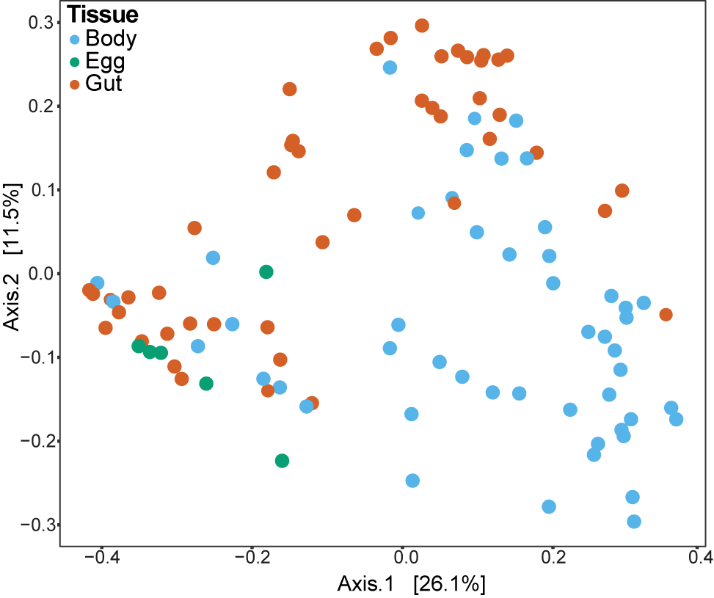

Supplemental Figure S6) PCoA plot of unweighted unifrac distances between microbial communities from *E. corrusca* adult guts, adult bodies, and eggs. Tissue types are indicated by color. n = 100

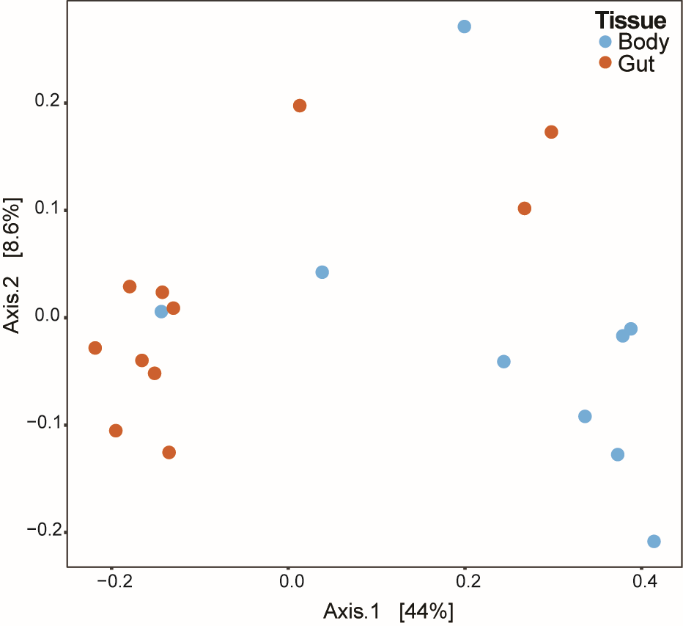

Supplemental Figure S7) PCoA plot of Unweighted Unifrac distances between the microbial communities from *Ph. versicolor* adult guts and bodies. Tissue type is indicated by color. n = 20

Table S1) PERMANOVA analyses testing the variation of beta-diversity explained by species and tissue type when comparing all 4 firefly taxa (*E. corrusca* adults & eggs, *Pyractomena borealis* larvae and whole adults, *Pyropyga decipiens* adults, and *Ph. versicolor* adults) n = 133

| Distance Metric | F. Model | R^2^ | Pr(>F) | Comparison |
| --- | --- | --- | --- | --- |
| Weighted Unifrac | 13.724 | 0.215 | 0.001*** | Taxon |
|  | 6.081 | 0.095 | 0.001*** | Tissue |
|  | 3.045 | 0.048 | 0.003** | Taxon:Tissue |
| Unweighted Unifrac | 9.014 | 0.157 | 0.001*** | Taxon |
|  | 5.560 | 0.097 | 0.001*** | Tissue |
|  | 1.692 | 0.030 | 0.022* | Taxon:Tissue |

Significance codes: ‘***’≥ 0.001, ‘**’≥ 0.01, ‘*’ ≥0.05

Table S2) PERMANOVA analyses testing the variation of beta-diversity explained by taxon and tissue type when comparing *E. corrusca* and *Ph. versicolor* adults. n = 114

| Distance Metric | F. Model | R^2^ | Pr(>F) | Comparison |
| --- | --- | --- | --- | --- |
| Weighted Unifrac | 23.626 | 0.164 | 0.001*** | Taxon |
|  | 1.994 | 0.014 | 0.105 | Tissue |
|  | 7.598 | 0.053 | 0.001*** | Taxon:Tissue |
| Unweighted Unifrac | 16.251 | 0.116 | 0.001*** | Taxon |
|  | 9.941 | 0.071 | 0.001*** | Tissue |
|  | 3.167 | 0.023 | 0.004** | Taxon:Tissue |

Significance codes: ‘***’≥ 0.001, ‘**’≥ 0.01

Table S3) PERMANOVA analyses testing the variation of beta-diversity explained by taxon, sex, and tissue type when comparing *Pyropyga* *decipiens*, *Ph. versicolor*, and *E. corrusca* for which sex was determined*.* n = 92

| Distance Metric | F. Model | R^2^ | Pr(>F) | Comparison |
| --- | --- | --- | --- | --- |
| Weighted Unifrac | 11.86 | 0.206 | 0.001*** | Taxon |
|  | 2.061 | 0.018 | 0.077 | Sex |
|  | 1.706 | 0.015 | 0.152 | Tissue |
|  | 0.557 | 0.010 | 0.827 | Taxon:Sex |
|  | 2.124 | 0.037 | 0.033* | Taxon:Tissue |
|  | 0.360 | 0.003 | 0.849 | Sex:Tissue |
|  | 0.717 | 0.006 | 0.527 | Taxon:Sex:Tissue |
| Unweighted Unifrac | 7.383 | 0.135 | 0.001*** | Taxon |
|  | 0.838 | 0.008 | 0.566 | Sex |
|  | 6.544 | 0.060 | 0.001*** | Tissue |
|  | 0.795 | 0.013 | 0.734 | Taxon:Sex |
|  | 1.403 | 0.026 | 0.088 | Taxon:Tissue |
|  | 1.083 | 0.010 | 0.324 | Sex:Tissue |
|  | 0.659 | 0.006 | 0.816 | Taxon:Sex:Tissue |

Significance codes: ‘***’≥ 0.001, ‘*’ ≥0.05

Table S4) PERMANOVA analyses testing the variation of beta-diversity explained by tissue type when comparing *E. corrusca* adults & eggs, *E. corrusca* adults, *Pyropyga decipiens*, *Ph. versicolor*, and *Pyractomena borealis* larvae & whole adults.

| **Species** | **Distance Metric** | **F. Model** | **R^2^** | **Pr(>F)** | **Comparison** |
| --- | --- | --- | --- | --- | --- |
| *E*. *corrusca* adults & eggs n = 101 | WUF | 6.834 | 0.122 | 0.001*** | Tissue |
|  | UUF | 6.856 | 0.123 | 0.001*** | Tissue |
| *E*. *corrusca* adults n = 96 | WUF | 0.805 | 0.009 | 0.542 | Tissue |
|  | UUF | 7.238 | 0.072 | 0.001*** | Tissue |
| *Ph. versicolor* adults n = 20 | WUF | 9.254 | 0.340 | 0.001*** | Tissue |
|  | UUF | 5.113 | 0.221 | 0.002*** | Tissue |
| *P. decipiens* adults n = 4 | WUF | 0.909 | 0.312 | 0.667 | Tissue |
|  | UUF | 1.220 | 0.379 | 0.333 | Tissue |
| *P.* *borealis* larvae & adult n = 8 | WUF | 2.429 | 0.493 | 0.060 | Tissue |
|  | UUF | 4.015 | 0.616 | 0.008** | Tissue |

Significance codes: ‘***’≥ 0.001, ‘**’≥ 0.01
