## Supplemental File2: Bioinformatic Code for "North American fireflies host low bacterial diversity"

^c^ Deceased

Emily A Green: https://orcid.org/0000-0002-0824-6369

Jonathan Klassen PhD: https://orcid.org/0000-0003-1745-8838

```
#Scripts used for firefly analysis
#Emily A. Green, University of Connecticut, Molecular and Cell Biology, Klassen lab

#Set working directory
#Load packages
library(dada2)
library(phyloseq)
library(ggplot2)
library(plyr)
library(vegan)
#Set theme, personal preference
theme_set(theme_bw())

#Dada2 pipeline followed with edits to the filtering/trimmming strictness
#Script is created to be run in an R environment, please check with dada2 for Windows & Linux specifications
#Set path to folder with sequence files.
path <- "#Path To Folder With Sequences"
#Read in your files, should match the file names
fnFs <- sort(list.files(path, pattern="_R1_001.fastq", full.name = TRUE))
fnRs <- sort(list.files(path, pattern="_R2_001.fastq", full.name = TRUE))
sample.names <- sapply(strsplit(basename(fnFs), "_"), `[`, 1)
#Check read quality
plotQualityProfile(fnFs[1:2])
plotQualityProfile(fnRs[1:2])
#Filter & trim reads into a specified folder
filt_path <- file.path(path, "filtered")
filtFs <- file.path(filt_path, paste0(sample.names, "_F_filt.fastq"))
filtRs <- file.path(filt_path, paste0(sample.names, "_R_filt.fastq"))
out <- filterAndTrim(fnFs, filtFs, fnRs, filtRs, truncLen = c(240,160), maxN=0, maxEE = c(2,5), truncQ=2, rm.phix =TRUE, compress=TRUE, multithread = TRUE)
#Check for any errors
errF <- learnErrors(filtFs, multithread = TRUE)
errR <- learnErrors(filtRs, multithread = TRUE)
plotErrors(errF, nominalQ=TRUE)
#Dereplicate sequences
derepFs <- derepFastq(filtFs, verbose=TRUE)
derepRs <- derepFastq(filtRs, verbose=TRUE)
names(derepFs) <- sample.names
names(derepRs) <- sample.names
#Create ASVs
dadaFs <- dada(derepFs, err=errF, multithread=TRUE)
dadaRs <- dada(derepRs, err=errR, multithread=TRUE)
dadaFs[[1]]
#Merge forward and reverse reads
mergers <- mergePairs(dadaFs, derepFs, dadaRs, derepRs, verbose = TRUE)
#Create Sequence table & remove chimeras
seqtab <- makeSequenceTable(mergers)
seqtab.nochim <- removeBimeraDenovo(seqtab, method="consensus", multithread=TRUE, verbose = TRUE)
sum(seqtab.nochim)/sum(seqtab)
#Track read count changes through current pipeline, can output a barplot of reads and a csv file.
getN <- function(x) sum(getUniques(x))
track <- cbind(out, sapply(dadaFs, getN), sapply(mergers, getN), rowSums(seqtab), rowSums(seqtab.nochim))
colnames(track) <- c("input", "filtered", "denoised", "merged", "tabled", "nonchim")
rownames(track) <- sample.names
barplot(t(track),beside=T,legend.text = T,las=2,cex.names=0.45,col=c("black","red","blue","yellow","purple","green"),space=c(0.2,2),ylim=c(0,120000),args.legend = list(cex=0.6))
write.csv(track, "TrackInfo.csv")
#Assign taxa names, use reference database of choice & include filepath.
taxa <- assignTaxonomy(seqtab.nochim, "silva_nr_v128_train_set.fa", multithread = TRUE)
taxa.print <- taxa
rownames(taxa.print) <- NULL
head(taxa.print)
#Create output file of all taxa names and sequences
write.csv(taxa.print, "TaxaList.csv")
#Create output file of all samples with taxa counts
write.csv(seqtab.nochim, "SampleASVCountList.csv")


#Read in Metadata from specified folder
meta <- read.csv("metadata.csv",header = TRUE, row.names = 1)
#create phyloseq object with metadata
ps <- phyloseq(otu_table(seqtab.nochim, taxa_are_rows = FALSE), sample_data(meta), tax_table(taxa))
ps

#load program
library(decontam)

#Check that metadata variables are present
sample_variables(ps)
#Decontam process, compares controls to samples for contaminates
df <- as.data.frame(sample_data(ps))
df$LibrarySize <- sample_sums(ps)
df <- df[order(df$LibrarySize),]
df$Index <- seq(nrow(df))
ggplot(data=df, aes(x=Index, y=LibrarySize, color=Sample_or_Control)) + geom_point()
sample_data(ps)$is.neg <- sample_data(ps)$Sample_or_Control == "Control Sample"
contamdf.prev <- isContaminant(ps, method="prevalence", neg="is.neg")
#Outputs TRUE or FALSE for contaminants present
table(contamdf.prev$contaminant)

#Create phyloseq objects
#Select sequences that are only in the Kingdom Bacteria
ps1 <- subset_taxa(ps,Kingdom == "Bacteria")
sample_data(ps1)
get_taxa_unique(ps1, taxonomic.rank = "Kingdom")
#Remove any sequence that is not identified at Phylum level
ps2 <- subset_taxa(ps1, !is.na(Phylum))
sort(get_taxa_unique(ps2, taxonomic.rank = "Phylum"))
#Remove Mitochondria sequences
ps3 <- subset_taxa(ps2, !Family == "Mitochondria")
sort(get_taxa_unique(ps3, taxonomic.rank = "Family"))
ps3
#Remove blanks and water control samples
ps4 <- subset_samples(ps3, !sample_data(ps3)$Sample_or_Control == "Control")
sample_data(ps4)

#Rarefy samples to and even depth
ps5 <- rarefy_even_depth(ps4, sample.size = 10000, rngseed = TRUE)

#Alpha Diversity done with samples with 10,000 reads but before Relative Abundance data
ps4.5 <- subset_samples(ps4, sample_sums(ps4)>= 10000)
#Only Firefly Dataset
ps4.5Fire <- subset_samples(ps4.5, !sample_data(ps4.5)$Species == "Tenebrio")
ps4.5Fire <- subset_samples(ps4.5Fire, !sample_data(ps4.5Fire)$Part == "Cotton")

#Violin Plot of alpha diversity
plot_richness(ps4.5Fire, x = "Species", measures = c("Shannon", "Simpson"), color = "Part")+ geom_violin() + theme(panel.background = element_blank(), panel.border = element_rect(fill = NA))
plot_richness(ps4.5Fire, x = "Species", measures = c("Shannon", "Simpson"), color = "Part")+ geom_violin() + theme(panel.background = element_blank(), panel.border = element_rect(fill = NA)) + stat_summary(fun.y= median, geom="point", shape=23, size=2)

#Estimate richness output, uses the alpha values in dataframe
FireflyAlphaDiv.df <- estimate_richness(ps4.5Fire, measures = c("Simpson", "Shannon"))
FireflyAlphaDiv.df$Part <- sample_data(ps4.5Fire)$Part
FireflyAlphaDiv.df$Species <- sample_data(ps4.5Fire)$Species

#Will run Kruskal-Wallis test, stats for if alpha div. differs between species or parts.
dunn.test(FireflyAlphaDiv.df$Shannon, g = FireflyAlphaDiv.df$Species, method = "bonferroni")
dunn.test(FireflyAlphaDiv.df$Simpson, g = FireflyAlphaDiv.df$Species, method = "bonferroni")
dunn.test(FireflyAlphaDiv.df$Shannon, g = FireflyAlphaDiv.df$Part, method = "bonferroni")
dunn.test(FireflyAlphaDiv.df$Simpson, g = FireflyAlphaDiv.df$Part, method = "bonferroni")

#Phyloseq object of only Fireflies (no Tenebrio)
psFire <- subset_samples(ps5, !sample_data(ps5)$Species == "Tenebrio")
psFire <- subset_samples(psFire, !sample_data(psFire)$Part == "Cotton")
psFire.glom <- tax_glom(psFire, taxrank = "Phylum")
psFire.glom.df <- psmelt(psFire.glom)
psFire.glom.df$Phylum <- as.character(psFire.glom.df$Phylum)
psFire.barplot.phylum <- ggplot(psFire.glom.df, aes(x = Sample, y = Abundance, fill = Phylum, scale_color_brewer(pallette = "Set3"))) + geom_bar(stat = "identity") + theme(axis.text.x = element_text(angle = 270, vjust = 0.5)) + facet_grid(~Species*Part, scales = "free", space = "free") + theme(strip.text = element_text(angle = 315), strip.background = element_blank(), panel.background = element_blank(), panel.grid = element_blank(), panel.border = element_blank())
plot(psFire.barplot.phylum)


#Phyloseq object of E. corrusca and P. borealis
psEllPb <- subset_samples(psFire, !sample_data(psFire)$Species == "Photuris")
psEllPb <- subset_samples(psEllPb, !sample_data(psEllPb)$Species == "Pyropyga")


#Phyloseq object of E. corrusca and Pyropyga
psEllPyr <- subset_samples(psFire, !sample_data(psFire)$Species == "Photuris")
psEllPyr <- subset_samples(psEllPyr, !sample_data(psEllPyr)$Species == "Pyractomena borealis")


#Phyloseq object of P. borealis and Pyropyga
psPbPyr <- subset_samples(psFire, !sample_data(psFire)$Species == "Photuris")
psPbPyr <- subset_samples(psPbPyr, !sample_data(psPbPyr)$Species == "Ellychnia corrusca")


#Phyloseq object of Photuris and P. borealis
psPbPho <- subset_samples(psFire, !sample_data(psFire)$Species == "Pyropyga")
psPbPho <- subset_samples(psPbPho, !sample_data(psPbPho)$Species == "Ellychnia corrusca")


#Phyloseq object of Photuris and Pyropyga
psPhoPyr <- subset_samples(psFire, !sample_data(psFire)$Species == "Pyractomena borealis")
psPhoPyr <- subset_samples(psPhoPyr, !sample_data(psPhoPyr)$Species == "Ellychnia corrusca")


#Phyloseq object of just Photuris
psPho <- subset_samples(ps5.rab, Species == "Photuris")
psPho.glom <- tax_glom(psPho, taxrank = "Phylum")
psPho.glom.df <- psmelt(psPho.glom)
psPho.glom.df$Phylum <- as.character(psPho.glom.df$Phylum)
theme_bw()
psPho.barplot.phylum <- ggplot(psPho.glom.df, aes(x = Sample, y = Abundance, fill = Phylum, scale_color_brewer(pallette = "Set3"))) + geom_bar(stat = "identity") + theme(axis.text.x = element_text(angle = 270, vjust = 0.5)) + facet_grid(~Part, scales = "free", space = "free") + theme(strip.text = element_text(angle = 315), strip.background = element_blank(), panel.background = element_blank(), panel.grid = element_blank(), panel.border = element_blank())
#psPho.barplot.phylum <- psPho.barplot.phylum + theme(panel.background = element_blank())
plot(psPho.barplot.phylum)
#15% other Phyla
max <- ddply(psPho.glom.df, ~Phylum, function(x) c(max = max(x$Abundance)))
Other <- max[max$max <= 0.15,]$Phylum
psPho.glom.df[psPho.glom.df$Phylum %in% Other,]$Phylum <- "Other"
psPho.barplot.phylum <- ggplot(psPho.glom.df, aes(x = Sample, y = Abundance, fill = Phylum, scale_color_brewer(pallette = "Set3"))) + geom_bar(stat = "identity") + theme(axis.text.x = element_text(angle = 270, vjust = 0.5))
plot(psPho.barplot.phylum)
#Arrange By Part
psPho.barplot.phylum <- ggplot(psPho.glom.df, aes(x = Sample, y = Abundance, fill = Phylum, scale_color_brewer(pallette = "Set3"))) + geom_bar(stat = "identity") + theme(axis.text.x = element_text(angle = 270, vjust = 0.5)) + facet_grid(~Part, scales = "free", space = "free") + theme(strip.text = element_text(angle = 315), strip.background = element_blank(), panel.background = element_blank() ,panel.grid = element_blank(), panel.border = element_blank())


#Output of specific phyla for Photuris
#Tenericutes
psPhoTen <- subset_taxa(psPho, Phylum == "Tenericutes")
psPhoTen.df <- psmelt(psPhoTen)
psPhoTen.df$Genus <- as.character(psPhoTen.df$Genus)
psPhoTen.barplot.phylum <- ggplot(psPhoTen.df, aes(x = Sample, y = Abundance, fill = Genus, scale_color_brewer(pallette = "Set3"))) + geom_bar(stat = "identity") + theme(axis.text.x = element_text(angle = 270, vjust = 0.5)) + facet_grid(~Part, scales = "free", space = "free") + theme(strip.text = element_text(angle = 315), strip.background = element_blank(), panel.background = element_blank(), panel.grid = element_blank(), panel.border = element_blank())
plot(psPhoTen.barplot.phylum)
#Proteobacteria
psPhoPro <- subset_taxa(psPho, Phylum == "Proteobacteria")
psPhoPro.df <- psmelt(psPhoPro)
psPhoPro.df$Genus <- as.character(psPhoPro.df$Genus)
max <- ddply(psPhoPro.df, ~Genus, function(x) c(max = max(x$Abundance)))
Other <- max[max$max <= 0.15,]$Genus
psPhoPro.df[psPhoPro.df$Genus %in% Other,]$Genus <- "Other"
psPhoPro.barplot.phylum <- ggplot(psPhoPro.df, aes(x = Sample, y = Abundance, fill = Genus, scale_color_brewer(pallette = "Set3"))) + geom_bar(stat = "identity") + theme(axis.text.x = element_text(angle = 270, vjust = 0.5)) + facet_grid(~Part, scales = "free", space = "free") + theme(strip.text = element_text(angle = 315), strip.background = element_blank(), panel.background = element_blank(), panel.grid = element_blank(), panel.border = element_blank())
plot(psPhoPro.barplot.phylum)
#Actinobacteria
psPhoAct <- subset_taxa(psPho, Phylum == "Actinobacteria")
psPhoAct.df <- psmelt(psPhoAct)
psPhoAct.df$Genus <- as.character(psPhoAct.df$Genus)
max <- ddply(psPhoAct.df, ~Genus, function(x) c(max = max(x$Abundance)))
Other <- max[max$max <= 0.15,]$Genus
psPhoAct.df[psPhoAct.df$Genus %in% Other,]$Genus <- "Other"
psPhoAct.barplot.phylum <- ggplot(psPhoAct.df, aes(x = Sample, y = Abundance, fill = Genus, scale_color_brewer(palette = "Set3"))) + geom_bar(stat = "identity") + theme(axis.text.x = element_text(angle = 270, vjust = 0.5)) + facet_grid(~Part, scales = "free", space = "free") + theme(strip.text = element_text(angle = 315), strip.background = element_blank(), panel.background = element_blank(), panel.grid = element_blank(), panel.border = element_blank())
plot(psPhoAct.barplot.phylum)
#Bacteroidetes
psPhoBac <- subset_taxa(psPho, Phylum == "Bacteroidetes")
psPhoBac.df <- psmelt(psPhoBac)
psPhoBac.df$Genus <- as.character(psPhoBac.df$Genus)
max <- ddply(psPhoBac.df, ~Genus, function(x) c(max = max(x$Abundance)))
Other <- max[max$max <= 0.15,]$Genus
psPhoBac.df[psPhoBac.df$Genus %in% Other,]$Genus <- "Other"
psPhoBac.barplot.phylum <- ggplot(psPhoBac.df, aes(x = Sample, y = Abundance, fill = Genus, scale_color_brewer(palette = "Set3"))) + geom_bar(stat = "identity") + theme(axis.text.x = element_text(angle = 270, vjust = 0.5)) + facet_grid(~Part, scales = "free", space = "free") + theme(strip.text = element_text(angle = 315), strip.background = element_blank(), panel.background = element_blank(), panel.grid = element_blank(), panel.border = element_blank())
plot(psPhoBac.barplot.phylum)


#Ellychnia corrusca Adults
psEllAd <- subset_samples(ps5.rab, Species == "Ellychnia corrusca")
psEllAd <- subset_samples(psEllAd, !sample_data(psEllAd)$Stage == "Egg")
psEllAd.glom <- tax_glom(psEllAd, taxrank = "Phylum")
psEllAd.glom.df <- psmelt(psEllAd.glom)
psEllAd.glom.df$Phylum <- as.character(psEllAd.glom.df$Phylum)
psEllAd.barplot.phylum <- ggplot(psEllAd.glom.df, aes(x = Sample, y = Abundance, fill = Phylum, scale_color_brewer(pallette = "Set3"))) + geom_bar(stat = "identity") + theme(axis.text.x = element_text(angle = 270, vjust = 0.5)) + facet_grid(~Part, scales = "free", space = "free") + theme(strip.text = element_text(angle = 315), strip.background = element_blank(), panel.background = element_blank(), panel.grid = element_blank(), panel.border = element_blank())
plot(psEllAd.barplot.phylum)
#15% other Phyla
max <- ddply(psEllAd.glom.df, ~Phylum, function(x) c(max = max(x$Abundance)))
Other <- max[max$max <= 0.15,]$Phylum
psEllAd.glom.df[psEllAd.glom.df$Phylum %in% Other,]$Phylum <- "Other"
psEllAd.barplot.phylum <- ggplot(psEllAd.glom.df, aes(x = Sample, y = Abundance, fill = Phylum, scale_color_brewer(pallette = "Set3"))) + geom_bar(stat = "identity") + theme(axis.text.x = element_text(angle = 270, vjust = 0.5)) + facet_grid(~Part, scales = "free", space = "free") + theme(strip.text = element_text(angle = 315), strip.background = element_blank(), panel.background = element_blank(), panel.grid = element_blank(), panel.border = element_blank())
plot(psEllAd.barplot.phylum)

#Output of specific phyla for E. corrusca
#Firmicutes
psEllAdFirm <- subset_taxa(psEllAd, Phylum == "Firmicutes")
psEllAdFirm.df <- psmelt(psEllAdFirm)
psEllAdFirm.df$Genus <- as.character(psEllAdFirm.df$Genus)
max <- ddply(psEllAdFirm.df, ~Genus, function(x) c(max = max(x$Abundance)))
Other <- max[max$max <= 0.15,]$Genus
psEllAdFirm.df[psEllAdFirm.df$Genus %in% Other,]$Genus <- "Other"
psEllAdFirm.barplot.phylum <- ggplot(psEllAdFirm.df, aes(x = Sample, y = Abundance, fill = Genus, scale_color_brewer(pallette = "Set3"))) + geom_bar(stat = "identity") + theme(axis.text.x = element_text(angle = 270, vjust = 0.5)) + facet_grid(~Stage, scales = "free", space = "free") + theme(strip.text = element_text(angle = 315), strip.background = element_blank(), panel.background = element_blank())+ facet_grid(~Part, scales = "free", space = "free") + theme(strip.text = element_text(angle = 315), strip.background = element_blank(), panel.background = element_blank(), panel.grid = element_blank(), panel.border = element_blank())
plot(psEllAdFirm.barplot.phylum)
#Tenericutes
psEllAdTen <- subset_taxa(psEllAd, Phylum == "Tenericutes")
psEllAdTen.df <- psmelt(psEllAdTen)
psEllAdTen.df$Phylum <- as.character(psEllAdTen.df$Phylum)
psEllAdTen.barplot.phylum <- ggplot(psEllAdTen.df, aes(x = Sample, y = Abundance, fill = Genus, scale_color_brewer(pallette = "Set3"))) + geom_bar(stat = "identity") + theme(axis.text.x = element_text(angle = 270, vjust = 0.5)) + facet_grid(~Part, scales = "free", space = "free") + theme(strip.text = element_text(angle = 315), strip.background = element_blank(), panel.background = element_blank(), panel.grid = element_blank(), panel.border = element_blank())
plot(psEllAdTen.barplot.phylum)
#Proteobacteria
# psEllAdPro.df$Genus[is.na(psEllAdPro.df$Genus)] <- "Not Assigned"
psEllAdPro <- subset_taxa(psEllAd, Phylum == "Proteobacteria")
psEllAdPro.df <- psmelt(psEllAdPro)
psEllAdPro.df$Genus <- as.character(psEllAdPro.df$Genus)
max <- ddply(psEllAdPro.df, ~Genus, function(x) c(max = max(x$Abundance)))
Other <- max[max$max <= 0.15,]$Genus
psEllAdPro.df[psEllAdPro.df$Genus %in% Other,]$Genus <- "Other"
psEllAdPro.barplot.phylum <- ggplot(psEllAdPro.df, aes(x = Sample, y = Abundance, fill = Genus, scale_color_brewer(pallette = "Set3"))) + geom_bar(stat = "identity") + theme(axis.text.x = element_text(angle = 270, vjust = 0.5)) + facet_grid(~Part, scales = "free", space = "free") + theme(strip.text = element_text(angle = 315), strip.background = element_blank(), panel.background = element_blank(), panel.grid = element_blank(), panel.border = element_blank())
plot(psEllAdPro.barplot.phylum)


#Ellychnia corrusca  eggs
psEllegg <- subset_samples(ps5.rab, Species == "Ellychnia corrusca")
psEllegg<- subset_samples(psEllegg, !sample_data(psEllegg)$Stage == "Adult")
psEllegg.glom <- tax_glom(psEllegg, taxrank = "Phylum")
psEllegg.glom.df <- psmelt(psEllegg.glom)
psEllegg.glom.df$Phylum <- as.character(psEllegg.glom.df$Phylum)
psEllegg.barplot.phylum <- ggplot(psEllegg.glom.df, aes(x = Sample, y = Abundance, fill = Phylum, scale_color_brewer(pallette = "Set3"))) + geom_bar(stat = "identity") + theme(axis.text.x = element_text(angle = 270, vjust = 0.5)) + facet_grid(~Part, scales = "free", space = "free") + theme(strip.text = element_text(angle = 315), strip.background = element_blank(), panel.background = element_blank(), panel.grid = element_blank(), panel.border = element_blank())
plot(psEllegg.barplot.phylum)
#15% other Phyla
max <- ddply(psEllegg.glom.df, ~Phylum, function(x) c(max = max(x$Abundance)))
Other <- max[max$max <= 0.15,]$Phylum
psEllegg.glom.df[psEllegg.glom.df$Phylum %in% Other,]$Phylum <- "Other"
psEllegg.barplot.phylum <- ggplot(psEllegg.glom.df, aes(x = Sample, y = Abundance, fill = Phylum, scale_color_brewer(pallette = "Set3"))) + geom_bar(stat = "identity") + theme(axis.text.x = element_text(angle = 270, vjust = 0.5)) + facet_grid(~Part, scales = "free", space = "free") + theme(strip.text = element_text(angle = 315), strip.background = element_blank(), panel.background = element_blank(), panel.grid = element_blank(), panel.border = element_blank())
plot(psEllegg.barplot.phylum)
#Remove Cotton Blanks
psEllegg2<- subset_samples(psEllegg, !sample_data(psEllegg)$Part == "Cotton")
psEllegg2.glom <- tax_glom(psEllegg2, taxrank = "Phylum")
psEllegg2.glom.df <- psmelt(psEllegg2.glom)
psEllegg2.glom.df$Phylum <- as.character(psEllegg2.glom.df$Phylum)
psEllegg2.barplot.phylum <- ggplot(psEllegg2.glom.df, aes(x = Sample, y = Abundance, fill = Phylum, scale_color_brewer(pallette = "Set3"))) + geom_bar(stat = "identity") + theme(axis.text.x = element_text(angle = 270, vjust = 0.5)) + facet_grid(~Part, scales = "free", space = "free") + theme(strip.text = element_text(angle = 315), strip.background = element_blank(), panel.background = element_blank(), panel.grid = element_blank(), panel.border = element_blank())
plot(psEllegg2.barplot.phylum)
#15% other Phyla
max <- ddply(psEllegg2.glom.df, ~Phylum, function(x) c(max = max(x$Abundance)))
Other <- max[max$max <= 0.15,]$Phylum
psEllegg2.glom.df[psEllegg2.glom.df$Phylum %in% Other,]$Phylum <- "Other"
psEllegg2.barplot.phylum


#Ellychnia egg Proteobacteria
#TO FIX THE NA ISSUES: psElleggPro.df$Genus[is.na(psElleggPro.df$Genus)] <- "Not Assigned"
psElleggPro <- subset_taxa(psEllegg2, Phylum == "Proteobacteria")
psElleggPro.df <- psmelt(psElleggPro)
psElleggPro.df$Genus <- as.character(psElleggPro.df$Genus)
max <- ddply(psElleggPro.df, ~Genus, function(x) c(max = max(x$Abundance)))
Other <- max[max$max <= 0.15,]$Genus
psElleggPro.df[psElleggPro.df$Genus %in% Other,]$Genus <- "Other"
psElleggPro.barplot.phylum <- ggplot(psElleggPro.df, aes(x = Sample, y = Abundance, fill = Genus, scale_color_brewer(pallette = "Set3"))) + geom_bar(stat = "identity") + theme(axis.text.x = element_text(angle = 270, vjust = 0.5)) + facet_grid(~Part, scales = "free", space = "free") + theme(strip.text = element_text(angle = 315), strip.background = element_blank(), panel.background = element_blank(), panel.grid = element_blank(), panel.border = element_blank())
plot(psElleggPro.barplot.phylum)
#Tenericutes
psEllegg2Ten<- subset_taxa(psEllegg2, Phylum == "Tenericutes")
psEllegg2Ten.df <- psmelt(psEllegg2Ten)
psEllegg2Ten.df$Genus <- as.character(psEllegg2Ten.df$Genus)
max <- ddply(psEllegg2Ten.df, ~Genus, function(x) c(max = max(x$Abundance)))
Other <- max[max$max <= 0.15,]$Genus
psEllegg2Ten.df[psEllegg2Ten.df$Genus %in% Other,]$Genus <- "Other"
psEllegg2Ten.barplot.phylum <- ggplot(psEllegg2Ten.df, aes(x = Sample, y = Abundance, fill = Genus, scale_color_brewer(pallette = "Set3"))) + geom_bar(stat = "identity") + theme(axis.text.x = element_text(angle = 270, vjust = 0.5)) + facet_grid(~Part, scales = "free", space = "free") + theme(strip.text = element_text(angle = 315), strip.background = element_blank(), panel.background = element_blank(), panel.grid = element_blank(), panel.border = element_blank())
plot(psEllegg2Ten.barplot.phylum)


#Phyloseq object of single P. borealis adult
psPbA <- subset_samples(ps5.rab, Species == "Pyractomena borealis")
psPbA<- subset_samples(psPbA, !sample_data(psPbA)$Stage == "Larvae")
psPbA.glom <- tax_glom(psPbA, taxrank = "Phylum")
psPbA.glom.df <- psmelt(psPbA.glom)
psPbA.glom.df$Phylum <- as.character(psPbA.glom.df$Phylum)
psPbA.barplot.phylum <- ggplot(psPbA.glom.df, aes(x = Sample, y = Abundance, fill = Phylum, scale_color_brewer(pallette = "Set3"))) + geom_bar(stat = "identity") + theme(axis.text.x = element_text(angle = 270, vjust = 0.5)) + facet_grid(~Part, scales = "free", space = "free") + theme(strip.text = element_text(angle = 315), strip.background = element_blank(), panel.background = element_blank(), panel.grid = element_blank(), panel.border = element_blank())
plot(psPbA.barplot.phylum)
#15% other Phyla
max <- ddply(psPbA.glom.df, ~Phylum, function(x) c(max = max(x$Abundance)))
Other <- max[max$max <= 0.15,]$Phylum
psPbA.glom.df[psPbA.glom.df$Phylum %in% Other,]$Phylum <- "Other"
psPbA.barplot.phylum <- ggplot(psPbA.glom.df, aes(x = Sample, y = Abundance, fill = Phylum, scale_color_brewer(pallette = "Set3"))) + geom_bar(stat = "identity") + theme(axis.text.x = element_text(angle = 270, vjust = 0.5)) + facet_grid(~Part, scales = "free", space = "free") + theme(strip.text = element_text(angle = 315), strip.background = element_blank(), panel.background = element_blank() ,panel.grid = element_blank(), panel.border = element_blank())
plot(psPbA.barplot.phylum)

#Phyloseq object of P. borealis larvae
psPbL <- subset_samples(ps5.rab, Species == "Pyractomena borealis")
psPbL <- subset_samples(psPbL, !sample_data(psPbL)$Stage == "Adult")
psPbL.glom <- tax_glom(psPbL, taxrank = "Phylum")
psPbL.glom.df <- psmelt(psPbL.glom)
psPbL.glom.df$Phylum <- as.character(psPbL.glom.df$Phylum)
psPbL.barplot.phylum <- ggplot(psPbL.glom.df, aes(x = Sample, y = Abundance, fill = Phylum, scale_color_brewer(pallette = "Set3"))) + geom_bar(stat = "identity") + theme(axis.text.x = element_text(angle = 270, vjust = 0.5)) + facet_grid(~Part, scales = "free", space = "free") + theme(strip.text = element_text(angle = 315), strip.background = element_blank(), panel.background = element_blank(), panel.grid = element_blank(), panel.border = element_blank())
plot(psPbL.barplot.phylum)
#15% other Phyla
max <- ddply(psPbL.glom.df, ~Phylum, function(x) c(max = max(x$Abundance)))
Other <- max[max$max <= 0.15,]$Phylum
psPbL.glom.df[psPbL.glom.df$Phylum %in% Other,]$Phylum <- "Other"
psPbL.barplot.phylum <- ggplot(psPbL.glom.df, aes(x = Sample, y = Abundance, fill = Phylum, scale_color_brewer(pallette = "Set3"))) + geom_bar(stat = "identity") + theme(axis.text.x = element_text(angle = 270, vjust = 0.5)) + facet_grid(~Part, scales = "free", space = "free") + theme(strip.text = element_text(angle = 315), strip.background = element_blank(), panel.background = element_blank() ,panel.grid = element_blank(), panel.border = element_blank())
plot(psPbL.barplot.phylum)

#Phyloseq object of all P. borealis
psPb <- subset_samples(ps5.rab, Species == "Pyractomena borealis")
psPb.glom <- tax_glom(psPb, taxrank = "Phylum")
psPb.glom.df <- psmelt(psPb.glom)
psPb.glom.df$Phylum <- as.character(psPb.glom.df$Phylum)
psPb.barplot.phylum <- ggplot(psPb.glom.df, aes(x = Sample, y = Abundance, fill = Phylum, scale_color_brewer(pallette = "Set3"))) + geom_bar(stat = "identity") + theme(axis.text.x = element_text(angle = 270, vjust = 0.5)) + facet_grid(~Part, scales = "free", space = "free") + theme(strip.text = element_text(angle = 315), strip.background = element_blank(), panel.background = element_blank(), panel.grid = element_blank(), panel.border = element_blank())
plot(psPb.barplot.phylum)
#15% other phyla
max <- ddply(psPb.glom.df, ~Phylum, function(x) c(max = max(x$Abundance)))
Other <- max[max$max <= 0.15,]$Phylum
psPb.glom.df[psPb.glom.df$Phylum %in% Other,]$Phylum <- "Other"
psPb.barplot.phylum <- ggplot(psPb.glom.df, aes(x = Sample, y = Abundance, fill = Phylum, scale_color_brewer(pallette = "Set3"))) + geom_bar(stat = "identity") + theme(axis.text.x = element_text(angle = 270, vjust = 0.5)) + facet_grid(~Part, scales = "free", space = "free") + theme(strip.text = element_text(angle = 315), strip.background = element_blank(), panel.background = element_blank() ,panel.grid = element_blank(), panel.border = element_blank())
plot(psPb.barplot.phylum)

#Output of specific phyla for P. borealis
#Proteobacteria
psPbPro <- subset_taxa(psPb, Phylum == "Proteobacteria")
psPbPro.df <- psmelt(psPbPro)
psPbPro.df$Genus <- as.character(psPbPro.df$Genus)
max <- ddply(psPbPro.df, ~Genus, function(x) c(max = max(x$Abundance)))
Other <- max[max$max <= 0.15,]$Genus
psPbPro.df[psPbPro.df$Genus %in% Other,]$Genus <- "Other"
psPbPro.barplot.phylum <- ggplot(psPbPro.df, aes(x = Sample, y = Abundance, fill = Genus, scale_color_brewer(pallette = "Set3"))) + geom_bar(stat = "identity") + theme(axis.text.x = element_text(angle = 270, vjust = 0.5)) + facet_grid(~Part, scales = "free", space = "free") + theme(strip.text = element_text(angle = 315), strip.background = element_blank(), panel.background = element_blank(), panel.grid = element_blank(), panel.border = element_blank())
plot(psPbPro.barplot.phylum)
#Actinobacteria
psPbAct <- subset_taxa(psPb, Phylum == "Actinobacteria")
psPbAct.df <- psmelt(psPbAct)
psPbAct.df$Genus <- as.character(psPbAct.df$Genus)
max <- ddply(psPbAct.df, ~Genus, function(x) c(max = max(x$Abundance)))
Other <- max[max$max <= 0.15,]$Genus
psPbAct.df[psPbAct.df$Genus %in% Other,]$Genus <- "Other"
psPbAct.barplot.phylum <- ggplot(psPbAct.df, aes(x = Sample, y = Abundance, fill = Genus, scale_color_brewer(pallette = "Set3"))) + geom_bar(stat = "identity") + theme(axis.text.x = element_text(angle = 270, vjust = 0.5)) + facet_grid(~Part, scales = "free", space = "free") + theme(strip.text = element_text(angle = 315), strip.background = element_blank(), panel.background = element_blank(), panel.grid = element_blank(), panel.border = element_blank())
plot(psPbAct.barplot.phylum)
#Tenericutes
psPbTen <- subset_taxa(psPb, Phylum == "Tenericutes")
psPbTen.df <- psmelt(psPbTen)
psPbTen.df$Genus <- as.character(psPbTen.df$Genus)
max <- ddply(psPbTen.df, ~Genus, function(x) c(max = max(x$Abundance)))
Other <- max[max$max <= 0.15,]$Genus
psPbTen.df[psPbTen.df$Genus %in% Other,]$Genus <- "Other"
psPbTen.barplot.phylum <- ggplot(psPbTen.df, aes(x = Sample, y = Abundance, fill = Genus, scale_color_brewer(pallette = "Set3"))) + geom_bar(stat = "identity") + theme(axis.text.x = element_text(angle = 270, vjust = 0.5)) + facet_grid(~Part, scales = "free", space = "free") + theme(strip.text = element_text(angle = 315), strip.background = element_blank(), panel.background = element_blank(), panel.grid = element_blank(), panel.border = element_blank())
plot(psPbTen.barplot.phylum)
#Bacteroidetes
psPbBac <- subset_taxa(psPb, Phylum == "Bacteroidetes")
psPbBac.df <- psmelt(psPbBac)
psPbBac.df$Genus <- as.character(psPbBac.df$Genus)
max <- ddply(psPbBac.df, ~Genus, function(x) c(max = max(x$Abundance)))
Other <- max[max$max <= 0.15,]$Genus
psPbBac.df[psPbBac.df$Genus %in% Other,]$Genus <- "Other"
psPbBac.barplot.phylum <- ggplot(psPbBac.df, aes(x = Sample, y = Abundance, fill = Genus, scale_color_brewer(pallette = "Set3"))) + geom_bar(stat = "identity") + theme(axis.text.x = element_text(angle = 270, vjust = 0.5)) + facet_grid(~Part, scales = "free", space = "free") + theme(strip.text = element_text(angle = 315), strip.background = element_blank(), panel.background = element_blank(), panel.grid = element_blank(), panel.border = element_blank())
plot(psPbBac.barplot.phylum)


#Phyloseq object of just Pyropyga
psPyr <- subset_samples(ps5.rab, Species == "Pyropyga")
psPyr.glom <- tax_glom(psPyr, taxrank = "Phylum")
psPyr.glom.df <- psmelt(psPyr.glom)
psPyr.glom.df$Phylum <- as.character(psPyr.glom.df$Phylum)
psPyr.barplot.phylum <- ggplot(psPyr.glom.df, aes(x = Sample, y = Abundance, fill = Phylum, scale_color_brewer(pallette = "Set3"))) + geom_bar(stat = "identity") + theme(axis.text.x = element_text(angle = 270, vjust = 0.5)) + facet_grid(~Part, scales = "free", space = "free") + theme(strip.text = element_text(angle = 315), strip.background = element_blank(), panel.background = element_blank(), panel.grid = element_blank(), panel.border = element_blank())
plot(psPyr.barplot.phylum)
#15% other phyla
max <- ddply(psPyr.glom.df, ~Phylum, function(x) c(max = max(x$Abundance)))
Other <- max[max$max <= 0.15,]$Phylum
psPyr.glom.df[psPyr.glom.df$Phylum %in% Other,]$Phylum <- "Other"
psPyr.barplot.phylum <- ggplot(psPyr.glom.df, aes(x = Sample, y = Abundance, fill = Phylum, scale_color_brewer(pallette = "Set3"))) + geom_bar(stat = "identity") + theme(axis.text.x = element_text(angle = 270, vjust = 0.5)) + facet_grid(~Part, scales = "free", space = "free") + theme(strip.text = element_text(angle = 315), strip.background = element_blank(), panel.background = element_blank() ,panel.grid = element_blank(), panel.border = element_blank())
plot(psPyr.barplot.phylum)

#Output of specific phyla for Pyropyga
#Tenericutes
psPyrTen<- subset_taxa(psPyr, Phylum == "Tenericutes")
psPyrTen.df <- psmelt(psPyrTen)
psPyrTen.df$Genus <- as.character(psPyrTen.df$Genus)
psPyrTen.barplot.phylum <- ggplot(psPyrTen.df, aes(x = Sample, y = Abundance, fill = Genus, scale_color_brewer(pallette = "Set3"))) + geom_bar(stat = "identity") + theme(axis.text.x = element_text(angle = 270, vjust = 0.5)) + facet_grid(~Part, scales = "free", space = "free") + theme(strip.text = element_text(angle = 315), strip.background = element_blank(), panel.background = element_blank(), panel.grid = element_blank(), panel.border = element_blank())
plot(psPyrTen.barplot.phylum)
#Proteobacteria
psPyrPro<- subset_taxa(psPyr, Phylum == "Proteobacteria")
psPyrPro.df <- psmelt(psPyrPro)
psPyrPro.df$Genus <- as.character(psPyrPro.df$Genus)
max <- ddply(psPyrPro.df, ~Genus, function(x) c(max = max(x$Abundance)))
Other <- max[max$max <= 0.15,]$Genus
psPyrPro.df[psPyrPro.df$Genus %in% Other,]$Genus <- "Other"
psPyrPro.barplot.phylum <- ggplot(psPyrPro.df, aes(x = Sample, y = Abundance, fill = Genus, scale_color_brewer(pallette = "Set3"))) + geom_bar(stat = "identity") + theme(axis.text.x = element_text(angle = 270, vjust = 0.5)) + facet_grid(~Part, scales = "free", space = "free") + theme(strip.text = element_text(angle = 315), strip.background = element_blank(), panel.background = element_blank(), panel.grid = element_blank(), panel.border = element_blank())
plot(psPyrPro.barplot.phylum)


#Compare Photuris and E. corrusca adults
#Phyloseq object of Photuris and E. corrusca adults
psPhoEll<- subset_samples(ps5.rab, !sample_data(ps5.rab)$Species == "Pyropyga")
psPhoEll<- subset_samples(psPhoEll, !sample_data(psPhoEll)$Species == "Pyractomena borealis")
psPhoEll<- subset_samples(psPhoEll, !sample_data(psPhoEll)$Species == "Pyropyga")
psPhoEll<- subset_samples(psPhoEll, !sample_data(psPhoEll)$Species == "Tenebrio")
psPhoEll<- subset_samples(psPhoEll, !sample_data(psPhoEll)$Part == "Egg")


#Load package
library(DESeq2)
#Making phyloseq object of raw data for each firefly species (not relative abundance) as DESeq2 requires non relative abundance data
#Photuris
psPhotnotrab <- subset_samples(ps5, sample_data(ps5)$Species == "Photuris")
#E. corrusca adults & eggs
psEllnotrab <- subset_samples(ps5, sample_data(ps5)$Species == "Ellychnia corrusca")
#E. corrusca adults
psEllAdnotrab <- subset_samples(psEllnotrab, !sample_data(psEllnotrab)$Stage == "Egg")
#Pyropyga
psPyrnotrab <- subset_samples(ps5, sample_data(ps5)$Species == "Pyropyga")
#P. borealis larvae and adult
psPbnotrab <- subset_samples(ps5, sample_data(ps5)$Species == "Pyractomena borealis")
#P. borealis laravae
psPbLnotrab <- subset_samples(psPbnotrab, !sample_data(psPbnotrab)$Stage == "Adult")


#Photuris DESeq2
psPhotnotrab <- subset_samples(ps5, sample_data(ps5)$Species == "Photuris")
psPhotnotrabdeseq = phyloseq_to_deseq2(psPhotnotrab, ~Part)
psPhotnotrabdeseq = DESeq(psPhotnotrabdeseq,  test="Wald", fitType = "parametric")
resPhot = results(psPhotnotrabdeseq, cooksCutoff = FALSE)
alpha = 0.01
sigtab_psPhotnotrabdeseq = resPhot[which(resPhot$padj < alpha), ]
sigtab_psPhotnotrabdeseq = cbind(as(sigtab_psPhotnotrabdeseq, "data.frame"), as(tax_table(psPhotnotrab)[rownames(sigtab_psPhotnotrabdeseq), ], "matrix"))
head(sigtab_psPhotnotrabdeseq)
dim(sigtab_psPhotnotrabdeseq)
scale_fill_discrete <- function(palname = "Set1", ...) {scale_fill_brewer(palette = palname, ...) }
# Phylum order
x = tapply(sigtab_psPhotnotrabdeseq$log2FoldChange, sigtab_psPhotnotrabdeseq$Phylum, function(x) max(x))
x = sort(x, TRUE)
sigtab_psPhotnotrabdeseq$Phylum = factor(as.character(sigtab_psPhotnotrabdeseq$Phylum), levels=names(x))
# Genus order
x = tapply(sigtab_psPhotnotrabdeseq$log2FoldChange, sigtab_psPhotnotrabdeseq$Genus, function(x) max(x))
x = sort(x, TRUE)
sigtab_psPhotnotrabdeseq$Genus = factor(as.character(sigtab_psPhotnotrabdeseq$Genus), levels=names(x))
ggplot(sigtab_psPhotnotrabdeseq, aes(x=Genus, y=log2FoldChange, color=Phylum)) + geom_point(size=6) + theme(axis.text.x = element_text(angle = -90, hjust = 0, vjust=0.5))

#E. corrusca DESeq2
psEllnotrab <- subset_samples(ps5, sample_data(ps5)$Species == "Ellychnia corrusca")
psEllnotrabdeseq = phyloseq_to_deseq2(psEllnotrab, ~Stage)
Ellgm_mean = function(x, na.rm=TRUE){exp(sum(log(x[x > 0]), na.rm=na.rm) / length(x))}
EllgeoMeans = apply(counts(psEllnotrabdeseq), 1, EllAdgm_mean)
psEllnotrabdeseq = estimateSizeFactors(psEllnotrabdeseq, geoMeans = geoMeans)
psEllnotrabdeseq = DESeq(psEllnotrabdeseq, fitType="local")
psEllnotrabdeseq = DESeq(psEllnotrabdeseq,  test="Wald", fitType = "parametric")
resEll = results(psEllnotrabdeseq, cooksCutoff = FALSE)
alpha = 0.01
sigtab_psEllnotrabdeseq = resEllAd[which(resEllAd$padj < alpha), ]
sigtab_psEllnotrabdeseq = cbind(as(sigtab_psEllAdnotrabdeseq, "data.frame"), as(tax_table(psEllAdnotrab)[rownames(sigtab_psEllAdnotrabdeseq), ], "matrix"))
head(sigtab_psEllAdnotrabdeseq)
dim(sigtab_psEllAdnotrabdeseq)
scale_fill_discrete <- function(palname = "Set1", ...) {scale_fill_brewer(palette = palname, ...) }
# Phylum order
x = tapply(sigtab_psPhoEllnotrabdeseq$log2FoldChange, sigtab_psEllnotrabdeseq$Phylum, function(x) max(x))
x = sort(x, TRUE)
sigtab_psEllnotrabdeseq$Phylum = factor(as.character(sigtab_psEllnotrabdeseq$Phylum), levels=names(x))
# Genus order
x = tapply(sigtab_psEllnotrabdeseq$log2FoldChange, sigtab_psEllnotrabdeseq$Genus, function(x) max(x))
x = sort(x, TRUE)
sigtab_psEllnotrabdeseq$Genus = factor(as.character(sigtab_psEllnotrabdeseq$Genus), levels=names(x))
ggplot(sigtab_psEllnotrabdeseq, aes(x=Genus, y=log2FoldChange, color=Phylum)) + geom_point(size=6) + theme(axis.text.x = element_text(angle = -90, hjust = 0, vjust=0.5))

#E. corrusca adults DESeq2
psEllnotrab <- subset_samples(ps5, sample_data(ps5)$Species == "Ellychnia corrusca")
psEllAdnotrab <- subset_samples(psEllnotrab, !sample_data(psEllnotrab)$Stage == "Egg")
psEllAdnotrabdeseq = phyloseq_to_deseq2(psEllAdnotrab, ~Part)
EllAdgm_mean = function(x, na.rm=TRUE){exp(sum(log(x[x > 0]), na.rm=na.rm) / length(x))}
EllAdgeoMeans = apply(counts(psEllAdnotrabdeseq), 1, EllAdgm_mean)
psEllAdnotrabdeseq = estimateSizeFactors(psEllAdnotrabdeseq, geoMeans = EllAdgeoMeans)
psEllAdnotrabdeseq = DESeq(psEllAdnotrabdeseq, fitType="local")
psEllAdnotrabdeseq = DESeq(psEllAdnotrabdeseq,  test="Wald", fitType = "parametric")
resEllAd = results(psEllAdnotrabdeseq, cooksCutoff = FALSE)
alpha = 0.01
sigtab_psEllAdnotrabdeseq = resEllAd[which(resEllAd$padj < alpha), ]
sigtab_psEllAdnotrabdeseq = cbind(as(sigtab_psEllAdnotrabdeseq, "data.frame"), as(tax_table(psEllAdnotrab)[rownames(sigtab_psEllAdnotrabdeseq), ], "matrix"))
head(sigtab_psEllAdnotrabdeseq)
dim(sigtab_psEllAdnotrabdeseq)
scale_fill_discrete <- function(palname = "Set1", ...) {scale_fill_brewer(palette = palname, ...) }
# Phylum order
x = tapply(sigtab_psEllAdnotrabdeseq$log2FoldChange, sigtab_psEllAdnotrabdeseq$Phylum, function(x) max(x))
x = sort(x, TRUE)
sigtab_psEllAdnotrabdeseq$Phylum = factor(as.character(sigtab_psEllAdnotrabdeseq$Phylum), levels=names(x))
# Genus order
x = tapply(sigtab_psEllAdnotrabdeseq$log2FoldChange, sigtab_psEllAdnotrabdeseq$Genus, function(x) max(x))
x = sort(x, TRUE)
sigtab_psEllAdnotrabdeseq$Genus = factor(as.character(sigtab_psEllAdnotrabdeseq$Genus), levels=names(x))
ggplot(sigtab_psEllAdnotrabdeseq, aes(x=Genus, y=log2FoldChange, color=Phylum)) + geom_point(size=6) + theme(axis.text.x = element_text(angle = -90, hjust = 0, vjust=0.5))

#DESeq2 species/species comparisons
#Photuris & E. corrusca
psPhoEllnotrab <- subset_samples(ps5, !sample_data(ps5)$Species == "Pyropyga")
psPhoEllnotrab <- subset_samples(psPhoEllnotrab, !sample_data(psPhoEllnotrab)$Species == "Pyractomena borealis")
psPhoEllnotrab <- subset_samples(psPhoEllnotrab, !sample_data(psPhoEllnotrab)$Species == "Pyropyga")
psPhoEllnotrab <- subset_samples(psPhoEllnotrab, !sample_data(psPhoEllnotrab)$Species == "Tenebrio")
psPhoEllnotrab <- subset_samples(psPhoEllnotrab, !sample_data(psPhoEllnotrab)$Part == "Egg")
psPhoEllnotrab <- subset_samples(psPhoEllnotrab, !sample_data(psPhoEllnotrab)$Part == "Cotton")
psPhoEllnotrabdeseq <- phyloseq_to_deseq2(psPhoEllnotrab, ~Species)
PhoEllgm_mean = function(x, na.rm=TRUE){exp(sum(log(x[x > 0]), na.rm=na.rm) / length(x))}
geoMeans = apply(counts(psPhoEllnotrabdeseq), 1, PhoEllgm_mean)
psPhoEllnotrabdeseq = estimateSizeFactors(psPhoEllnotrabdeseq, geoMeans = geoMeans)
psPhoEllnotrabdeseq = DESeq(psPhoEllnotrabdeseq, fitType="local")
psPhoEllnotrabdeseq = DESeq(psPhoEllnotrabdeseq,  test="Wald", fitType = "parametric")
resPhoEll = results(psPhoEllnotrabdeseq, cooksCutoff = FALSE)
alpha = 0.01
sigtab_psPhoEllnotrabdeseq = resPhoEll[which(resPhoEll$padj < alpha), ]
sigtab_psPhoEllnotrabdeseq = cbind(as(sigtab_psPhoEllnotrabdeseq, "data.frame"), as(tax_table(psPhoEllnotrab)[rownames(sigtab_psPhoEllnotrabdeseq), ], "matrix"))
head(sigtab_psPhoEllnotrabdeseq)
dim(sigtab_psPhoEllnotrabdeseq)
scale_fill_discrete <- function(palname = "Set1", ...) {scale_fill_brewer(palette = palname, ...) }
# Phylum order
x = tapply(sigtab_psPhoEllnotrabdeseq$log2FoldChange, sigtab_psPhoEllnotrabdeseq$Phylum, function(x) max(x))
x = sort(x, TRUE)
sigtab_psPhoEllnotrabdeseq$Phylum = factor(as.character(sigtab_psPhoEllnotrabdeseq$Phylum), levels=names(x))
# Genus order
x = tapply(sigtab_psPhoEllnotrabdeseq$log2FoldChange, sigtab_psPhoEllnotrabdeseq$Genus, function(x) max(x))
x = sort(x, TRUE)
sigtab_psPhoEllnotrabdeseq$Genus = factor(as.character(sigtab_psPhoEllnotrabdeseq$Genus), levels=names(x))
ggplot(sigtab_psPhoEllnotrabdeseq, aes(x=Genus, y=log2FoldChange, color=Phylum)) + geom_point(size=6) + theme(axis.text.x = element_text(angle = -90, hjust = 0, vjust=0.5))
#output data file
write.csv(sigtab_psPhoEllnotrabdeseq, "DeSeq Photuris Ellychnia.csv")


#Weighted and Unweighted Unifrac testing
#After uniquesToFasta command, align seqs of the fasta file and then create phylogenetic tree. upload BestTree into read_tree command. For this publication I used CIPRES (available at https://www.phylo.org/portal2/login!input.action). Sequences were aligned using MAFFT on XSEDE (7.402), and phylogenetic tree made using RAxML-HPC BlackBox (8.2.12).

#Only Photuris and E. corrusca Adult samples
testspecies <- c("Photuris", "Ellychnia corrusca")
psPEadults <- subset_samples(ps5, sample_data(ps5)$Species %in% testspecies)
psPEadults <- subset_samples(psPEadults, !sample_data(psPEadults)$Part == "Egg")
psPEadults <- subset_samples(psPEadults, !sample_data(psPEadults)$Stage == "Egg")
psPEadults.df <- t(data.frame(otu_table(psPEadults)))
psPEadults.df <- data.frame(psPEadults.df)
psPEadults.df$abundance <- rowSums(psPEadults.df[,c(1:115)])
psPEadults.df$sequence <- rownames(psPEadults.df)
psPEadults.uniques <- getUniques(psPEadults.df)
uniquesToFasta(psPEadults.uniques, "PEadults.fasta", ids = psPEadults.df$sequence)
#Make Tree files
PEadults.tree <- read_tree("Best Tree Result file")
psPEadults.tree <- phyloseq(otu_table(otu_table(psPEadults)),sample_data(sample_data(psPEadults)), tax_table(tax_table(psPEadults)), phy_tree(PEadults.tree))
#Unweighted unifrac
PEadults.uuf.dist <- distance(psPEadults.tree, method = "unifrac")
PEadults.uuf.ord <- ordinate(psPEadults.tree, method = "PCoA", distance = "unifrac")
PEadults.uuf.plot <- plot_ordination(psPEadults.tree, PEadults.uuf.ord, color = "Part", shape = "Species")
plot(PEadults.uuf.plot)
#PERMANOVA
psPEadults.tree.data <- data.frame(sample_data(psPEadults.tree))
adonis(PEadults.uuf.dist ~Species, psPEadults.tree.data)
adonis(PEadults.uuf.dist ~Part, psPEadults.tree.data)
adonis(PEadults.uuf.dist ~Species*Part, psPEadults.tree.data)
#Weighed Unifrac
PEadults.wuf.dist <- distance(psPEadults.tree, method = "wunifrac")
PEadults.wuf.ord <- ordinate(psPEadults.tree, method = "PCoA", distance = "wunifrac")
PEadults.wuf.plot <- plot_ordination(psPEadults.tree, PEadults.wuf.ord, color = "Species", shape = "Part")
plot(PEadults.wuf.plot)
#PERMANOVA
psPEadults.tree.data <- data.frame(sample_data(psPEadults.tree))
adonis(PEadults.wuf.dist ~Species, psPEadults.tree.data)
adonis(PEadults.wuf.dist ~Part, psPEadults.tree.data)
adonis(PEadults.wuf.dist ~Species*Part, psPEadults.tree.data)


#Photuris and E.corrusca including eggs
psPE <- subset_samples(ps5, sample_data(ps5)$Species %in% testspecies)
psPE <- subset_samples(psPE, !sample_data(psPE)$Part == "Cotton")
psPE.df <- t(data.frame(otu_table(psPE)))
psPE.df <- data.frame(psPE.df)
psPE.df$abundance <- rowSums(psPE.df[,c(1:121)])
psPE.df$sequence <- rownames(psPE.df)
psPE.uniques <- getUniques(psPEadults.df)
uniquesToFasta(psPE.uniques, "PE.fasta", ids = psPE.df$sequence)
#Make Tree files
PE.tree <- read_tree("Best Tree Result file")
psPE.tree <- phyloseq(otu_table(otu_table(psPE)),sample_data(sample_data(psPE)), tax_table(tax_table(psPE)), phy_tree(PE.tree))
#Unweighted unifrac
PE.uuf.dist <- distance(psPE.tree, method = "unifrac")
PE.uuf.ord <- ordinate(psPE.tree, method = "PCoA", distance = "unifrac")
PE.uuf.plot <- plot_ordination(psPE.tree, PE.uuf.ord, color = "Species", shape = "Part")
plot(PE.uuf.plot)
#PERMANOVA
psPE.tree.data <- data.frame(sample_data(psPE.tree))
adonis(PE.uuf.dist ~Species, psPE.tree.data)
adonis(PE.uuf.dist ~Part, psPE.tree.data)
adonis(PE.uuf.dist ~Species*Part, psPE.tree.data)
#Weighed Unifrac
PE.wuf.dist <- distance(psPE.tree, method = "wunifrac")
PE.wuf.ord <- ordinate(psPE.tree, method = "PCoA", distance = "wunifrac")
PE.wuf.plot <- plot_ordination(psPE.tree, PE.wuf.ord, color = "Part", shape = "Species")
plot(PE.wuf.plot)
#PERMANOVA
psPE.tree.data <- data.frame(sample_data(psPE.tree))
adonis(PE.wuf.dist ~Species, psPE.tree.data)
adonis(PE.wuf.dist ~Part, psPE.tree.data)
adonis(PE.wuf.dist ~Species*Part, psPE.tree.data)


#All Fireflies
psFire.df <- t(data.frame(otu_table(psFire)))
psFire.df <- data.frame(psFire.df)
psFire.df$abundance <- rowSums(psFire.df[,c(1:133)])
psFire.df$sequence <- rownames(psFire.df)
psFire.uniques <- getUniques(psFire.df)
uniquesToFasta(psFire.uniques, "Fire.fasta", ids = psFire.df$sequence)
#Make Tree files
Fire.tree <- read_tree("Best Tree Result file")
psFire.tree <- phyloseq(otu_table(otu_table(psFire)),sample_data(sample_data(psFire)), tax_table(tax_table(psFire)), phy_tree(Fire.tree))
#Unweighted unifrac
Fire.uuf.dist <- distance(psFire.tree, method = "unifrac")
Fire.uuf.ord <- ordinate(psFire.tree, method = "PCoA", distance = "unifrac")
Fire.uuf.plot <- plot_ordination(psFire.tree, Fire.uuf.ord, color = "Species", shape = "Part")
plot(Fire.uuf.plot)
#PERMANOVA
psFire.tree.data <- data.frame(sample_data(psFire.tree))
adonis(Fire.uuf.dist ~Species, psFire.tree.data)
adonis(Fire.uuf.dist ~Part, psFire.tree.data)
adonis(Fire.uuf.dist ~Species * Part, psFire.tree.data)
#Weighed Unifrac
Fire.wuf.dist <- distance(psFire.tree, method = "wunifrac")
Fire.wuf.ord <- ordinate(psFire.tree, method = "PCoA", distance = "wunifrac")
Fire.wuf.plot <- plot_ordination(psFire.tree, Fire.wuf.ord, color = "Species", shape = "Part")
plot(Fire.wuf.plot)
#PERMANOVA
psFire.tree.data <- data.frame(sample_data(psFire.tree))
adonis(Fire.wuf.dist ~Species, psFire.tree.data)
adonis(Fire.wuf.dist ~Part, psFire.tree.data)
adonis(Fire.wuf.dist ~Species * Part, psFire.tree.data)


#Single Species Weighted & Unweighted Unifrac tests
#E.corrusca adults + eggs
psEll2 <- psEll
psEll2.df <- t(data.frame(otu_table(psEll2)))
psEll2.df <- data.frame(psEll2.df)
psEll2.df$abundance <- rowSums(psEll2.df[,c(1:101)])
psEll2.df$sequence <- rownames(psEll2.df)
psEll2.uniques <- getUniques(psEll2.df)
uniquesToFasta(psEll2.uniques, "psEll2.fasta", ids = psEll2.df$sequence)
#Make Tree files
Ell2.tree <- read_tree("Best Tree Result file")
psEll2.tree <- phyloseq(otu_table(otu_table(psEll2)),sample_data(sample_data(psEll2)), tax_table(tax_table(psEll2)), phy_tree(Ell2.tree))
#Unweighted unifrac
Ell2.uuf.dist <- distance(psEll2.tree, method = "unifrac")
Ell2.uuf.ord <- ordinate(psEll2.tree, method = "PCoA", distance = "unifrac")
Ell2.uuf.plot <- plot_ordination(psEll2.tree, Ell2.uuf.ord, color = "Part")
plot(Ell2.uuf.plot)
#PERMANOVA
psEll2.tree.data <- data.frame(sample_data(psEll2.tree))
adonis(Ell2.uuf.dist ~Part, psEll2.tree.data)
#Weighed Unifrac
Ell2.wuf.dist <- distance(psEll2.tree, method = "wunifrac")
Ell2.wuf.ord <- ordinate(psEll2.tree, method = "PCoA", distance = "wunifrac")
Ell2.wuf.plot <- plot_ordination(psEll2.tree, Ell2.wuf.ord, color = "Part")
plot(Ell2.wuf.plot)
#PERMANOVA
psEll2.tree.data <- data.frame(sample_data(psEll2.tree))
adonis(Ell2.wuf.dist ~Part, psEll2.tree.data)

#E. corrusca adults only
psEllAd2 <- psEllAd
psEllAd2.df <- t(data.frame(otu_table(psEllAd2)))
psEllAd2.df <- data.frame(psEllAd2.df)
psEllAd2.df$abundance <- rowSums(psEllAd2.df[,c(1:95)])
psEllAd2.df$sequence <- rownames(psEllAd2.df)
psEllAd2.uniques <- getUniques(psEllAd2.df)
uniquesToFasta(psEllAd2.uniques, "psEllAd2.fasta", ids = psEllAd2.df$sequence)
#Make Tree files
EllAd2.tree <- read_tree("Best Tree Result file")
psEllAd2.tree <- phyloseq(otu_table(otu_table(psEllAd2)),sample_data(sample_data(psEllAd2)), tax_table(tax_table(psEllAd2)), phy_tree(EllAd2.tree))
#Unweighted unifrac
EllAd2.uuf.dist <- distance(psEllAd2.tree, method = "unifrac")
EllAd2.uuf.ord <- ordinate(psEllAd2.tree, method = "PCoA", distance = "unifrac")
EllAd2.uuf.plot <- plot_ordination(psEllAd2.tree, EllAd2.uuf.ord, color = "Part")
plot(EllAd2.uuf.plot)
#PERMANOVA
psEllAd2.tree.data <- data.frame(sample_data(psEllAd2.tree))
adonis(EllAd2.uuf.dist ~Part, psEllAd2.tree.data)
#Weighed Unifrac
EllAd2.wuf.dist <- distance(psEllAd2.tree, method = "wunifrac")
EllAd2.wuf.ord <- ordinate(psEllAd2.tree, method = "PCoA", distance = "wunifrac")
EllAd2.wuf.plot <- plot_ordination(psEllAd2.tree, EllAd2.wuf.ord, color = "Part")
plot(EllAd2.wuf.plot)
#PERMANOVA
psEllAd2.tree.data <- data.frame(sample_data(psEllAd2.tree))
adonis(EllAd2.wuf.dist ~Part, psEllAd2.tree.data)

#Photuris Adults
psPho2 <- psPho
psPho2.df <- t(data.frame(otu_table(psPho2)))
psPho2.df <- data.frame(psPho2.df)
psPho2.df$abundance <- rowSums(psPho2.df[,c(1:20)])
psPho2.df$sequence <- rownames(psPho2.df)
psPho2.uniques <- getUniques(psPho2.df)
uniquesToFasta(psPho2.uniques, "psPho2.fasta", ids = psPho2.df$sequence)
#Make Tree files
Pho2.tree <- read_tree("Best Tree Result file")
psPho2.tree <- phyloseq(otu_table(otu_table(psPho2)),sample_data(sample_data(psPho2)), tax_table(tax_table(psPho2)), phy_tree(Pho2.tree))
#Unweighted unifrac
Pho2.uuf.dist <- distance(psPho2.tree, method = "unifrac")
Pho2.uuf.ord <- ordinate(psPho2.tree, method = "PCoA", distance = "unifrac")
Pho2.uuf.plot <- plot_ordination(psPho2.tree, Pho2.uuf.ord, color = "Part")
plot(Pho2.uuf.plot)
#PERMANOVA
psPho2.tree.data <- data.frame(sample_data(psPho2.tree))
adonis(Pho2.uuf.dist ~Part, psPho2.tree.data)
#Weighed Unifrac
Pho2.wuf.dist <- distance(psPho2.tree, method = "wunifrac")
Pho2.wuf.ord <- ordinate(psPho2.tree, method = "PCoA", distance = "wunifrac")
Pho2.wuf.plot <- plot_ordination(psPho2.tree, Pho2.wuf.ord, color = "Part")
plot(Pho2.wuf.plot)
#PERMANOVA
psPho2.tree.data <- data.frame(sample_data(psPho2.tree))
adonis(Pho2.wuf.dist ~Part, psPho2.tree.data)

#P.borealis
psPb2 <- psPb
psPb2.df <- t(data.frame(otu_table(psPb2)))
psPb2.df <- data.frame(psPb2.df)
psPb2.df$abundance <- rowSums(psPb2.df[,c(1:8)])
psPb2.df$sequence <- rownames(psPb2.df)
psPb2.uniques <- getUniques(psPb2.df)
uniquesToFasta(psPb2.uniques, "psPb2.fasta", ids = psPb2.df$sequence)
#Make Tree files
Pb2.tree <- read_tree("#Best Tree Result file")
psPb2.tree <- phyloseq(otu_table(otu_table(psPb2)),sample_data(sample_data(psPb2)), tax_table(tax_table(psPb2)), phy_tree(Pb2.tree))
#Unweighted unifrac
Pb2.uuf.dist <- distance(psPb2.tree, method = "unifrac")
Pb2.uuf.ord <- ordinate(psPb2.tree, method = "PCoA", distance = "unifrac")
Pb2.uuf.plot <- plot_ordination(psPb2.tree, Pb2.uuf.ord, color = "Part")
plot(Pb2.uuf.plot)
#PERMANOVA
psPb2.tree.data <- data.frame(sample_data(psPb2.tree))
adonis(Pb2.uuf.dist ~Part, psPb2.tree.data)
#Weighed Unifrac
Pb2.wuf.dist <- distance(psPb2.tree, method = "wunifrac")
Pb2.wuf.ord <- ordinate(psPb2.tree, method = "PCoA", distance = "wunifrac")
Pb2.wuf.plot <- plot_ordination(psPb2.tree, Pb2.wuf.ord, color = "Part")
plot(Pb2.wuf.plot)
#PERMANOVA
psPb2.tree.data <- data.frame(sample_data(psPb2.tree))
adonis(Pb2.wuf.dist ~Part, psPb2.tree.data)

#Pyropyga
psPyr2 <- psPyr
psPyr2.df <- t(data.frame(otu_table(psPyr2)))
psPyr2.df <- data.frame(psPyr2.df)
psPyr2.df$abundance <- rowSums(psPyr2.df[,c(1:4)])
psPyr2.df$sequence <- rownames(psPyr2.df)
psPyr2.uniques <- getUniques(psPyr2.df)
uniquesToFasta(psPyr2.uniques, "psPyr2.fasta", ids = psPyr2.df$sequence)
#Make Tree files
Pyr2.tree <- read_tree("#Best Tree Result file")
psPyr2.tree <- phyloseq(otu_table(otu_table(psPyr2)),sample_data(sample_data(psPyr2)), tax_table(tax_table(psPyr2)), phy_tree(Pyr2.tree))
#Unweighted unifrac
Pyr2.uuf.dist <- distance(psPyr2.tree, method = "unifrac")
Pyr2.uuf.ord <- ordinate(psPyr2.tree, method = "PCoA", distance = "unifrac")
Pyr2.uuf.plot <- plot_ordination(psPyr2.tree, Pyr2.uuf.ord, color = "Part")
plot(Pyr2.uuf.plot)
#PERMANOVA
psPyr2.tree.data <- data.frame(sample_data(psPyr2.tree))
adonis(Pyr2.uuf.dist ~Part, psPyr2.tree.data)
#Weighed Unifrac
Pyr2.wuf.dist <- distance(psPyr2.tree, method = "wunifrac")
Pyr2.wuf.ord <- ordinate(psPyr2.tree, method = "PCoA", distance = "wunifrac")
Pyr2.wuf.plot <- plot_ordination(psPyr2.tree, Pyr2.wuf.ord, color = "Part")
plot(Pyr2.wuf.plot)
#PERMANOVA
psPyr2.tree.data <- data.frame(sample_data(psPyr2.tree))
adonis(Pyr2.wuf.dist ~Part, psPyr2.tree.data)


#Firefly combinations
#E. corrusca and Pyropyga
psFire.tree <- phyloseq(otu_table(otu_table(psFire)),sample_data(sample_data(psFire)), tax_table(tax_table(psFire)), phy_tree(Fire.tree))
psEllPyr.tree <- subset_samples(psFire.tree, !sample_data(psFire.tree)$Species == "Photuris")
psEllPyr.tree <- subset_samples(psEllPyr.tree, !sample_data(psEllPyr.tree)$Species == "Pyractomena borealis")
#Unweighted unifrac
EllPyr.uuf.dist <- distance(psEllPyr.tree, method = "unifrac")
EllPyr.uuf.ord <- ordinate(psEllPyr.tree, method = "PCoA", distance = "unifrac")
EllPyr.uuf.plot <- plot_ordination(psEllPyr.tree, EllPyr.uuf.ord, color = "Species", shape = "Part")
plot(EllPyr.uuf.plot)
#PERMANOVA
psEllPyr.tree.data <- data.frame(sample_data(psEllPyr.tree))
adonis(EllPyr.uuf.dist ~Species, psEllPyr.tree.data)
adonis(EllPyr.uuf.dist ~Part, psEllPyr.tree.data)
adonis(EllPyr.uuf.dist ~Species * Part, psEllPyr.tree.data)
#Weighted Unifrac
EllPyr.wuf.dist <- distance(psEllPyr.tree, method = "wunifrac")
EllPyr.wuf.ord <- ordinate(psEllPyr.tree, method = "PCoA", distance = "wunifrac")
EllPyr.wuf.plot <- plot_ordination(psEllPyr.tree, EllPyr.wuf.ord, color = "Species", shape = "Part")
plot(EllPyr.wuf.plot)
#PERMANOVA
psEllPyr.tree.data <- data.frame(sample_data(psEllPyr.tree))
adonis(EllPyr.wuf.dist ~Species, psEllPyr.tree.data)
adonis(EllPyr.wuf.dist ~Part, psEllPyr.tree.data)
adonis(EllPyr.wuf.dist ~Species * Part, psEllPyr.tree.data)

#E. corrusca and P. borealis
psEllPb.tree <- subset_samples(psFire.tree, !sample_data(psFire.tree)$Species == "Photuris")
psEllPb.tree <- subset_samples(psEllPb.tree, !sample_data(psEllPb.tree)$Species == "Pyropyga")
#Unweighted unifrac
EllPb.uuf.dist <- distance(psEllPb.tree, method = "unifrac")
EllPb.uuf.ord <- ordinate(psEllPb.tree, method = "PCoA", distance = "unifrac")
EllPb.uuf.plot <- plot_ordination(psEllPb.tree, EllPb.uuf.ord, color = "Species", shape = "Part")
plot(EllPb.uuf.plot)
#PERMANOVA
psEllPb.tree.data <- data.frame(sample_data(psEllPb.tree))
adonis(EllPb.uuf.dist ~Species, psEllPb.tree.data)
adonis(EllPb.uuf.dist ~Part, psEllPb.tree.data)
adonis(EllPb.uuf.dist ~Species * Part, psEllPb.tree.data)
#Weighted Unifrac
EllPb.wuf.dist <- distance(psEllPb.tree, method = "wunifrac")
EllPb.wuf.ord <- ordinate(psEllPb.tree, method = "PCoA", distance = "wunifrac")
EllPb.wuf.plot <- plot_ordination(psEllPb.tree, EllPb.wuf.ord, color = "Species", shape = "Part")
plot(EllPb.wuf.plot)
#PERMANOVA
psEllPb.tree.data <- data.frame(sample_data(psEllPb.tree))
adonis(EllPb.wuf.dist ~Species, psEllPb.tree.data)
adonis(EllPb.wuf.dist ~Part, psEllPb.tree.data)
adonis(EllPb.wuf.dist ~Species * Part, psEllPb.tree.data)

#Photuris and P. borealis
psPhoPb.tree <- subset_samples(psFire.tree, !sample_data(psFire.tree)$Species == "Ellychnia corrusca")
psPhoPb.tree <- subset_samples(psPhoPb.tree, !sample_data(psPhoPb.tree)$Species == "Pyropyga")
#Unweighted unifrac
PhoPb.uuf.dist <- distance(psPhoPb.tree, method = "unifrac")
PhoPb.uuf.ord <- ordinate(psPhoPb.tree, method = "PCoA", distance = "unifrac")
PhoPb.uuf.plot <- plot_ordination(psPhoPb.tree, PhoPb.uuf.ord, color = "Species", shape = "Part")
plot(PhoPb.uuf.plot)
#PERMANOVA
psPhoPb.tree.data <- data.frame(sample_data(psPhoPb.tree))
adonis(PhoPb.uuf.dist ~Species, psPhoPb.tree.data)
adonis(PhoPb.uuf.dist ~Part, psPhoPb.tree.data)
adonis(PhoPb.uuf.dist ~Species * Part, psPhoPb.tree.data)
#Weighted Unifrac
PhoPb.wuf.dist <- distance(psPhoPb.tree, method = "wunifrac")
PhoPb.wuf.ord <- ordinate(psPhoPb.tree, method = "PCoA", distance = "wunifrac")
PhoPb.wuf.plot <- plot_ordination(psPhoPb.tree, PhoPb.wuf.ord, color = "Species", shape = "Part")
plot(PhoPb.wuf.plot)
#PERMANOVA
psPhoPb.tree.data <- data.frame(sample_data(psPhoPb.tree))
adonis(PhoPb.wuf.dist ~Species, psPhoPb.tree.data)
adonis(PhoPb.wuf.dist ~Part, psPhoPb.tree.data)
adonis(PhoPb.wuf.dist ~Species * Part, psPhoPb.tree.data)

#Pyropyga and P. borealis
psPyrPb.tree <- subset_samples(psFire.tree, !sample_data(psFire.tree)$Species == "Ellychnia corrusca")
psPyrPb.tree <- subset_samples(psPyrPb.tree, !sample_data(psPyrPb.tree)$Species == "Photuris")
#Unweighted unifrac
PyrPb.uuf.dist <- distance(psPyrPb.tree, method = "unifrac")
PyrPb.uuf.ord <- ordinate(psPyrPb.tree, method = "PCoA", distance = "unifrac")
PyrPb.uuf.plot <- plot_ordination(psPyrPb.tree, PyrPb.uuf.ord, color = "Species", shape = "Part")
plot(PyrPb.uuf.plot)
#PERMANOVA
psPyrPb.tree.data <- data.frame(sample_data(psPyrPb.tree))
adonis(PyrPb.uuf.dist ~Species, psPyrPb.tree.data)
adonis(PyrPb.uuf.dist ~Part, psPyrPb.tree.data)
adonis(PyrPb.uuf.dist ~Species * Part, psPyrPb.tree.data)
#Weighted Unifrac
PyrPb.wuf.dist <- distance(psPyrPb.tree, method = "wunifrac")
PyrPb.wuf.ord <- ordinate(psPyrPb.tree, method = "PCoA", distance = "wunifrac")
PyrPb.wuf.plot <- plot_ordination(psPyrPb.tree, PyrPb.wuf.ord, color = "Species", shape = "Part")
plot(PyrPb.wuf.plot)
#PERMANOVA
psPyrPb.tree.data <- data.frame(sample_data(psPyrPb.tree))
adonis(PyrPb.wuf.dist ~Species, psPyrPb.tree.data)
adonis(PyrPb.wuf.dist ~Part, psPyrPb.tree.data)
adonis(PyrPb.wuf.dist ~Species * Part, psPyrPb.tree.data)

#Pyropyga and Photuris
psPyrPho.tree <- subset_samples(psFire.tree, !sample_data(psFire.tree)$Species == "Ellychnia corrusca")
psPyrPho.tree <- subset_samples(psPyrPho.tree, !sample_data(psPyrPho.tree)$Species == "Pyractomena borealis")
#Unweighted unifrac
PyrPho.uuf.dist <- distance(psPyrPho.tree, method = "unifrac")
PyrPho.uuf.ord <- ordinate(psPyrPho.tree, method = "PCoA", distance = "unifrac")
PyrPho.uuf.plot <- plot_ordination(psPyrPho.tree, PyrPho.uuf.ord, color = "Species", shape = "Part")
plot(PyrPho.uuf.plot)
#PERMANOVA
psPyrPho.tree.data <- data.frame(sample_data(psPyrPho.tree))
adonis(PyrPho.uuf.dist ~Species, psPyrPho.tree.data)
adonis(PyrPho.uuf.dist ~Part, psPyrPho.tree.data)
adonis(PyrPho.uuf.dist ~Species* Part, psPyrPho.tree.data)
#Weighted Unifrac
PyrPho.wuf.dist <- distance(psPyrPho.tree, method = "wunifrac")
PyrPho.wuf.ord <- ordinate(psPyrPho.tree, method = "PCoA", distance = "wunifrac")
PyrPho.wuf.plot <- plot_ordination(psPyrPho.tree, PyrPho.wuf.ord, color = "Species", shape = "Part")
plot(PyrPho.wuf.plot)
#PERMANOVA
psPyrPho.tree.data <- data.frame(sample_data(psPyrPho.tree))
adonis(PyrPho.wuf.dist ~Species, psPyrPho.tree.data)
adonis(PyrPho.wuf.dist ~Part, psPyrPho.tree.data)
adonis(PyrPho.wuf.dist ~Species * Part, psPyrPho.tree.data)

#Create a dataset of samples that have sex type meta data
psSex <- subset_samples(psFire, !sample_data(psFire)$Sex == "None")
psSex.rab <- transform_sample_counts(psSex, function(x) x/sum(x))
psSex.rab.glom <- tax_glom(psSex.rab, taxrank = "Phylum")
psSex.rab.glom.df <- psmelt(psSex.rab.glom)
psSex.rab.glom.df$Phylum <- as.character(psSex.rab.glom.df$Phylum)
psSex.barplot.phylum <- ggplot(psSex.rab.glom.df, aes(x = Sample, y = Abundance, fill = Phylum, scale_color_brewer(pallette = "Set3"))) + geom_bar(stat = "identity") + theme(axis.text.x = element_text(angle = 270, vjust = 0.5)) + facet_grid(~Species*Sex, scales = "free", space = "free") + theme(strip.text = element_text(angle = 315), strip.background = element_blank(), panel.background = element_blank())
plot(psSex.barplot.phylum)
#Top Phyla
max <- ddply(psSex.rab.glom.df, ~Phylum, function(x) c(max = max(x$Abundance)))
Other <- max[max$max <= 0.15,]$Phylum
psSex.rab.glom.df[psSex.rab.glom.df$Phylum %in% Other,]$Phylum <- "Other"


#Weighted & Unweighted Unifrac for all fireflies with sex type metadata
psSex.df <- t(data.frame(otu_table(psSex)))
psSex.df <- data.frame(psSex.df)
psSex.df$abundance <- rowSums(psSex.df[,c(1:92)])
psSex.df$sequence <- rownames(psSex.df)
psSex.uniques <- getUniques(psSex.df)
uniquesToFasta(psSex.uniques, "Sex.fasta", ids = psSex.df$sequence)
#Make Tree files
Sex.tree <- read_tree("#Best Tree Result file")
psSex.tree <- phyloseq(otu_table(otu_table(psSex)),sample_data(sample_data(psSex)), tax_table(tax_table(psSex)), phy_tree(Sex.tree))
#Unweighted unifrac
Sex.uuf.dist <- distance(psSex.tree, method = "unifrac")
Sex.uuf.ord <- ordinate(psSex.tree, method = "PCoA", distance = "unifrac")
Sex.uuf.plot <- plot_ordination(psSex.tree, Sex.uuf.ord, color = "Species", shape = "Sex")
plot(Sex.uuf.plot)
#PERMANOVA
psSex.tree.data <- data.frame(sample_data(psSex.tree))
adonis(Sex.uuf.dist ~Species * Sex, psSex.tree.data)

#Weighed Unifrac
Sex.wuf.dist <- distance(psSex.tree, method = "wunifrac")
Sex.wuf.ord <- ordinate(psSex.tree, method = "PCoA", distance = "wunifrac")
Sex.wuf.plot <- plot_ordination(psSex.tree, Sex.wuf.ord, color = "Species", shape = "Sex")
plot(Sex.wuf.plot)
#PERMANOVA
psSex.tree.data <- data.frame(sample_data(psSex.tree))
adonis(Sex.wuf.dist ~Species * Sex, psSex.tree.data)


#Just E. corrusca sex type data
psEcSex <- subset_samples(psSex, !sample_data(psSex)$Species == "Photuris")
psEcSex <- subset_samples(psEcSex, !sample_data(psEcSex)$Species == "Pyropyga")
psEcSex.df <- t(data.frame(otu_table(psEcSex)))
psEcSex.df <- data.frame(psEcSex.df)
psEcSex.df$abundance <- rowSums(psEcSex.df[,c(1:68)])
psEcSex.df$sequence <- rownames(psEcSex.df)
psEcSex.uniques <- getUniques(psEcSex.df)
uniquesToFasta(psEcSex.uniques, "EcSex.fasta", ids = psEcSex.df$sequence)
#Make Tree files
EcSex.tree <- read_tree("#Best Tree Result file")
psEcSex.tree <- phyloseq(otu_table(otu_table(psEcSex)),sample_data(sample_data(psEcSex)), tax_table(tax_table(psEcSex)), phy_tree(EcSex.tree))
#Unweighted unifrac
EcSex.uuf.dist <- distance(psEcSex.tree, method = "unifrac")
EcSex.uuf.ord <- ordinate(psEcSex.tree, method = "PCoA", distance = "unifrac")
EcSex.uuf.plot <- plot_ordination(psEcSex.tree, EcSex.uuf.ord, color = "Species", shape = "Sex")
plot(EcSex.uuf.plot)
#PERMANOVA
psEcSex.tree.data <- data.frame(sample_data(psEcSex.tree))
adonis(EcSex.uuf.dist ~Sex , psEcSex.tree.data)
#Weighed Unifrac
EcSex.wuf.dist <- distance(psEcSex.tree, method = "wunifrac")
EcSex.wuf.ord <- ordinate(psEcSex.tree, method = "PCoA", distance = "wunifrac")
EcSex.wuf.plot <- plot_ordination(psEcSex.tree, EcSex.wuf.ord, color = "Species", shape = "Sex")
plot(EcSex.wuf.plot)
#PERMANOVA
psEcSex.tree.data <- data.frame(sample_data(psEcSex.tree))
adonis(EcSex.wuf.dist ~Sex * Part, psEcSex.tree.data)

#Photuris sex type data
psPhoSex <- subset_samples(psSex, !sample_data(psSex)$Species == "Ellychnia corrusca")
psPhoSex <- subset_samples(psPhoSex, !sample_data(psPhoSex)$Species == "Pyropyga")
psPhoSex.df <- t(data.frame(otu_table(psPhoSex)))
psPhoSex.df <- data.frame(psPhoSex.df)
psPhoSex.df$abundance <- rowSums(psPhoSex.df[,c(1:20)])
psPhoSex.df$sequence <- rownames(psPhoSex.df)
psPhoSex.uniques <- getUniques(psPhoSex.df)
uniquesToFasta(psPhoSex.uniques, "PhoSex.fasta", ids = psPhoSex.df$sequence)
#Make Tree files
PhoSex.tree <- read_tree("#Best Tree Result file")
psPhoSex.tree <- phyloseq(otu_table(otu_table(psPhoSex)),sample_data(sample_data(psPhoSex)), tax_table(tax_table(psPhoSex)), phy_tree(PhoSex.tree))
#Unweighted unifrac
PhoSex.uuf.dist <- distance(psPhoSex.tree, method = "unifrac")
PhoSex.uuf.ord <- ordinate(psPhoSex.tree, method = "PCoA", distance = "unifrac")
PhoSex.uuf.plot <- plot_ordination(psPhoSex.tree, PhoSex.uuf.ord, color = "Species", shape = "Sex")
plot(PhoSex.uuf.plot)
#PERMANOVA
psPhoSex.tree.data <- data.frame(sample_data(psPhoSex.tree))
adonis(PhoSex.uuf.dist ~Part * Sex, psPhoSex.tree.data)
#Weighed Unifrac
PhoSex.wuf.dist <- distance(psPhoSex.tree, method = "wunifrac")
PhoSex.wuf.ord <- ordinate(psPhoSex.tree, method = "PCoA", distance = "wunifrac")
PhoSex.wuf.plot <- plot_ordination(psPhoSex.tree, PhoSex.wuf.ord, color = "Species", shape = "Sex")
plot(PhoSex.wuf.plot)
#PERMANOVA
psPhoSex.tree.data <- data.frame(sample_data(psPhoSex.tree))
adonis(PhoSex.wuf.dist ~Part * Sex, psPhoSex.tree.data)


#Heat map 11-13-19
library(gplots)
#Color Spectrum
colors = rev(brewer.pal(11, "Spectral"))
colors = colorRampPalette(colors)(50)
#Color palette
Bluecolor = colorRampPalette(colors = c("white","royalblue3"))

#Use Color palette at the end
heatmap.2(as.matrix(madtop25), margins = c(4,9), density.info = "none", trace = "none",cexCol = 0.3, distfun = function(x) dist(x, method = "euclidean"), hclustfun = function(x) hclust(x, method="ward.D"), col=colors)

#Use tax glom to get rid of ASV variants
psFire.glom.gen <- tax_glom(psFire, taxrank = "Genus")
psFire.glom.gen.rab <- transform_sample_counts(psFire.glom.gen, function(x) x/sum(x))
psFire.glom.gen.rab.df <- psmelt(psFire.glom.gen.rab)
#Use the psFire.glom.gen.rab.df and export the OTU Table as a csv for next portion.
#Use the otu table and taxa output of glom.gen.rab
hmdata3 <- read.csv("Retry Firefly data/GlomOTUTable.csv", header=TRUE, row.names = 1, sep = ",")
hmmeta <- read.csv("Retry Firefly data/MetaDataHeatMap.csv", header=TRUE,row.names=1, sep=",")
hmdata3 <- as.matrix(hmdata3[,-dim(hmdata3)[2]])

#Abundance data arranged in decreasing value
mad2 <- apply(hmdata3,1,sum)
mad2 <- sort(mad2, decreasing = T)

#Only include the top 26 bacteria genera
top26 <- mad2[1:26]
madtop26 <- hmdata3[row.names(hmdata3) %in% names(top26),]
#Euclidean
heatmap.2(as.matrix(madtop26), margins = c(4,9), density.info = "none", trace = "none",cexCol = 0.3, cexRow = 0.5, distfun = function(x) dist(x, method = "euclidean"), hclustfun = function(x) hclust(x, method="ward.D"), col=colors, Colv = FALSE, dendrogram = "row")


# Speic-Easi
library(SpiecEasi)
library(phyloseq)
library(Matrix)
library(igraph)

#Try to make a phyloseq object that does not have the asv's
#change asv sequence into a count
#se.OTU <- otu_table(psFire.glom.gen.rab)
#se.Tax <- tax_table(psFire.glom.gen.rab)
#se.Samples <- sample_data(psFire.glom.gen.rab)
#Need to make the asv seqs into a number

#made a dupicate of my ps object to mess around with
psFire.se <- psFire.glom.gen.rab
#make ASV names into a count
taxa_names(psFire.se) <- paste0("", seq(ntaxa(psFire.se)))

#Make a matrix with the same taxa as the heatmap! 2-13-2020
psFire.se.26 <- names(sort(taxa_sums(psFire.se), decreasing = TRUE))[1:26]
psFire.se.26top <- transform_sample_counts(psFire.se, function(x) x/sum(x))
psFire.se.26top <- prune_taxa(psFire.se.26, psFire.se.26top)

se.mb.ps26top <- spiec.easi(psFire.se.26top, method='mb', lambda.min.ratio=1e-2, nlambda=20, pulsar.params=list(rep.num=50))
ig6.mb <- adj2igraph(getRefit(se.mb.ps26top), vertex.attr=list(name=taxa_names(psFire.se.26top)))
sbeta_graph.ig6 <- symBeta(getOptBeta(se.mb.ps26top), mode = 'maxabs')
#Need to make weight into a vector where the actual weights are in column 3. found on github....causes error without
weights.ig6 <- Matrix::summary(t(sbeta_graph.ig6))[,3]
ig6.mb <- adj2igraph(getRefit(se.mb.ps26top), diag = TRUE, edge.attr= list(weight = weights.ig6), vertex.attr = list(name = taxa_names(psFire.se.26top)))
#export into cytoscape
library(RCy3)
createNetworkFromIgraph(ig6.mb, "Feb13Top26TaxaIg6")


#Phylogenetic Tree code
#2-19-2020
#The tree includes seqs found from fireflies, and a few others from Silva for tree rooting, and top mollicute ASVs from the data set.  TopFfMOlicuteASVs.fasta includes, Mesoplasma ASVs 1-4, 1 Entomoplasma ASV and Spiroplasma ASVs 1&2. Refined5SilvaMollicutes.fasta includes silva seqs of Spiroplasmas, Entomoplasmas, Mesoplasmas and rooting sequences. The KnownFfMollicutes.fasta has non-silva seqs that were isolated from fireflies.

cat TopFfMollicuteASVs.fasta Refined5SilvaMollicutes.fasta KnownFfMollicutes.fasta > Ref5FfMol.fasta
muscle -in Ref5FfMol.fasta -out AlRef_5FfMol.fasta
perl idk.phy AlRef_5FfMol.fasta
raxmlHPC -mGTRGAMMAI -s AlRef_5FfMol.fasta.phy -n Ref5FfMolTree -f a -N 500 -x 2 -p 5
```
